# An Observational Study of Ballooning in Large Spiders: Nanoscale Multi-Fibers Enable Large Spiders’ Soaring Flight

**DOI:** 10.1101/206334

**Authors:** Moonsung Cho, Peter Neubauer, Christoph Fahrenson, Ingo Rechenberg

## Abstract

The physical mechanism of aerial dispersal of spiders, “ballooning behavior,” is still unclear because of the lack of serious scientific observations and experiments. Therefore, as a first step in clarifying the phenomenon, we studied the ballooning behavior of relatively large spiders (heavier than 5 mg) in nature. Additional wind tunnel tests to identify ballooning silks were implemented in the laboratory. From our observation, it seems obvious that spiders actively evaluate the condition of the wind with their front leg (leg I) and wait for the preferable wind condition for their ballooning takeoff. In the wind tunnel tests, as yet unknown physical properties of ballooning fibers (length, thickness and number of fibers) were identified. Large spiders, 16–20 mg *Xysticus* species, spun 50 to 60 nanoscale fibers, with a diameter of 121 to 323 nm. The length of these threads was 3.22 ± 1.31 m (N = 22). These physical properties of ballooning fibers can explain the ballooning of large spiders with relatively light updrafts, 0.1–0.5 m s^-1^, which exist in a light breeze of 1.5–3.3 m s^-1^. Additionally, in line with previous research on turbulence in atmospheric boundary layers and from our wind measurements, it is hypothesized that spiders use the ascending air current for their aerial dispersal, the “ejection” regime, which is induced by hairpin vortices in the atmospheric boundary layer turbulence. This regime is highly correlated with lower wind speeds. This coincides well with the fact that spiders usually balloon when the wind speed is lower than 3 m s^-1^.

## 1. Introduction

Some spiders from different families, such as Linyphiidae (sheet-weaver spiders), Araneidae (orb-weaving spiders), Lycosidae (wolf spiders) and Thomisidae (crab spiders) can disperse aerially with the help of their silks, which is usually called ballooning behavior [1–6]. There are two representative takeoff methods in ballooning flight; “tiptoe” and “rafting” [7–10]. If spiders perceive appropriate weather conditions for ballooning, they climb up to the highest position of a blade of grass or a branch of a tree and raise their abdomen as if standing on their tiptoes, in order to position the abdomen at the highest level, before spinning the ballooning lines. They release a single or a number of silks in the wind current and wait until a sufficient updraft draws their body up in the air. This is known as a “tiptoe” takeoff [9,10] (see S1, S2). Another takeoff method is called “rafting,” where spiders release the ballooning lines from a hanging position relying on their drag line [7,8,10] (see S3). In this way, some spiders can travel passively hundreds of kilometers and can reach as high as 4.5 kilometers above sea level [11,12]. For example, one of the first immigrant species on new-born volcanic islands are known to be spiders [13-15]. Aerial dispersal of spiders is an influential factor on agricultural economy and ecology, because spiders are highly ranked predators in arthropods and impact on a prey’s population [16]. Due to the spider’s incredible aerial dispersal ability, the physical mechanism of a spider’s flight has been questioned for a long time, not only in public media but also in scientific research [16-23].

Ballooning dispersal is efficiently used by spiderlings (young spiders) just a few days after eclosion from their eggs) to avoid cannibalism at their birth sites, which are densely populated by hundreds of young spiders, and to reduce competition for resources [23,24]. Some adult female spiders balloon to find a place for a new colony [4,25,26] and others balloon to search for food and mates [4,27]. Most of the ballooning spiders were spiderlings and spiders under 3 mm in length and 0.2 to 2 mg in mass [1–5,28,29]. Nevertheless, there are only a few reports on the ballooning of large spiders (over 3 mm in length, over 5 mg in mass) [4,5,25,26].

Spiders balloon most frequently during late spring and autumn seasons [2,4,30]. The influences of microclimates on ballooning, such as temperature, humidity and wind conditions, have been extensively studied: (i) Many studies agree on a positive correlation of temperature [1,3,31] or a rapid increase in temperature [31–34]; (ii) low humidity is favorable for spiders to balloon [1,32,34]; (iii) for small spiders, 0.2–2 mm in length, the favorable mean wind speed is limited to 3 m s^-1^ at a level of 2 m [30,31,35]. The local favorable wind speeds were 0.35–1.7 m s^-1^ in experiments and 0.55–0.75 m s^-1^ in nature [1,3]. These values, however, differ for spiders of different sizes (between 0.78–1.21 mm) [1,36]. Recently, Lee et al. showed that not only the mean wind speed at a level of 2 m but also the local wind speed can be limited by a wind speed of 3 m s^-1^ for spiderlings [37]. Instability of atmosphere was pointed out as an influential factor [30,31,35]. Suter and Reynolds suggested a possible relation of spiders’ ballooning behavior with atmospheric turbulent flow [16,20].

There have been a number of models that have tried to explain spiders’ high buoyant capability (aerial dispersal capability): a fluid-dynamic lollipop model [17], a flexible filament model in turbulence [16] and an electrostatic flight model [22]. Recently, Zhao et al. implemented the two-dimensional numerical simulation using an immersed boundary method, which can simulate the ballooning dynamics in more detail [38]. The result shows that the atmospheric instability enables longer suspension of a ballooner in the air, which agrees with the result of Reynolds’s simulation and suggests that a spider may sense the vibration of vortex shedding on the spider silk through their silk [16,38], which is an interesting hypothesis.

In spite of the abovementioned models and studies, dynamics in spiders ballooning are still not well understood, because of a lack of serious scientific observation studies and specific experiments. Many of the ballooning spiders are very small with weights of 0.2–2 mg, which are difficult to study [1,3,36,37]. Many described experiments were not focusing on the spider’s ballooning behavior itself, but assumed that spiders use a certain length of the drag line [18,19,21]. The ballooning of large spiders is also a struggle because of (i) the observed physical properties of ballooning silks and spider size (60–80 cm long and 3–4 silk threads, 85 to 150 mg body weight) of an adult of *Stegodyphus mimosarum* seemed to be unrealistic for ballooning [25], because the required vertical speed of wind was 9.2 to 21.6 m s^-1^ according to Henschel’s calculation [18,39]; (ii) Humphrey’s model cannot explain the ballooning flight of spiders with a weight of over 9 mg, because of the mechanical properties of a spider silk [17]. The following questions are still to be answered: (i) How many and how long are the silk fibers needed for ballooning, especially in the case of large spiders with weights over 5 mg? (ii) Which silk fibers and glands are used for ballooning? (iii) How do ballooning silks shape during the flight? (iv) Do spiders control the buoyant capability by changing the length of silks or their pose during the flight? (v) Why do spiders usually balloon at a low wind speed (below 3 m s^-1^)?

The aim of this paper is to offer behavioral clues and quantitative data in ballooning flight that may answer these questions. Therefore, we investigated the ballooning behavior of adult and subadult crab spiders (*Xysticus* species, Thomisidae), that had a size of 3–6 mm and a weight of 6–25 mg. This observation of large spiders could provide a good basis for the physical characterization of ballooning. Additional experiments were performed in a wind tunnel, for a precise documentation of ballooning silks and to analyze the details of ballooning behavior. Also, the aerodynamic environment on a flat grass field was measured to investigate the usable updraft for a ballooning flight.

## 2. Materials and methods

### 2.1 Field observations

Crab spiders *(Xysticus cristatus, Xysticus audax*, etc.) were collected at Lilienthal Park and along the Teltow canal in the Berlin area, Germany, and observed each autumn, especially during October, from 2014 to 2016. Lilienthal Park was selected for the observation of pre-ballooning behavior, because the ballooning phenomenon of adult and subadult crab spiders is frequently observed in this region.

#### 2.1.1 Takeoff

On sunny and partly cloudy days in autumn, 14 crab spiders (8 females, 2 males and 4 not identified; adult or subadults) were collected at Lilienthal Park. These spiders were released in the same place on a self-built artificial mushroom-like platform (a 5.5 cm diameter half sphere, 1.2 m above a ground surface, gypsum material). This platform was intended to stimulate tiptoe pre-ballooning behavior. Because of its convex surface (half sphere), spiders can easily recognize that the top of the convex surface is the highest position that may promote tiptoe behavior. The white color of this platform allowed the visual clarity of the spider’s behavior. During the observation, the ballooning behaviors were recorded by digital camera. Additionally, titanium tetrachloride was used for the flow visualization of the wind. The local wind speed and temperature were not measured directly, but the values from the Dahlem weather station (4.5 km distance from the observation site, the anemometer is installed 36 m above the ground in the 20 m-high forest canopy) may provide approximate conditions.

#### 2.1.2 Statistical analysis of pre-ballooning behaviors

The 14 crab spiders were set a total of 25 times onto the mushroom shaped platform in natural weather. The detailed pre-ballooning behaviors in “takeoff” observation were analyzed statistically: (i) active sensing motion, (ii) tiptoe motion, (iii) tiptoe takeoff, (iv) drop down, (v) rafting takeoff (see “takeoff” in the results section). If they did not tend to balloon or hid for about 5 min, they were recollected. Once a spider raised one or both front legs (leg I) and then put them back again on the platform, this was considered to be an active sensing motion and was counted as one behavior. Tiptoe motion, raising the abdomen and putting it down again was also counted as one motion. The transitions between behaviors were counted. These numbers were first expressed as percentage frequency by normalizing with the total number of transitions (see equation (1), *f*_*i,j*_ is a normalized frequency of the transition from i-th to j-th behavior. *n*_*i,j*_ is the number of the transition from i-th to j-th. *m* is the number of categorized behaviors). The probability from one behavior to next behavior is expressed by normalizing with the total number of previous behaviors (see equation (2), *p*_*i,j*_ is the probability of the transition from i-th to j-th behavior.).

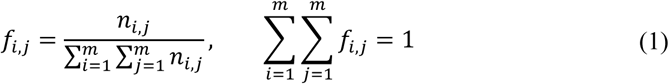

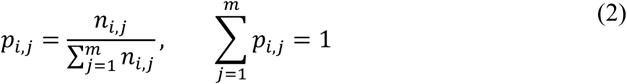

#### 2.1.3 Gliding

Gliding of crab spiders was observed in the autumn along the Teltow canal, due to the ecological and topographical benefits for the observation of ballooners: (i) crab spiders frequently glide along the canal during the autumn; (ii) the angle of the sun’s rays in the morning was appropriate for the detection of the ballooning silks during their flight; (iii) the dense trees on the opposite side of the canal provide a dark background, which facilitates the observation of floating silks. The shape of the threads during flight could not be photographed nor recorded as a video, because only limited parts of the ballooning threads were reflected by the sun’s rays. However, the movement and shape of the threads could be recognized with the naked eye and these shapes were quickly sketched by hand.

### 2.2 Identification of ballooning lines

#### 2.2.1 Sampling of ballooning lines

Twelve crab spiders (9 females and 3 males, adult or subadult) were used for the wind tunnel experiment. Pre-ballooning behavior of these spiders was induced in front of an open jet wind tunnel in which the diameter of the nozzle exit is 0.6 m (see Fig. 1A). The spinning behaviors that led to the ballooning silks was observed precisely. There were no obstacles next to the wind tunnel, leaving about 9 m of free space from the nozzle, to allow ballooning fibers to float horizontally without any adhesion to other objects. The wind speed and temperature were measured with a PL-135 HAN hot-wire anemometer. To enable sampling of the ballooning fibers, the wind speed was adjusted at 0.9 m s^-1^, because spiders drifted downstream if the wind speed was above 1 m s^-1^. The room temperature was 22°–25°C. The ballooning behavior was stimulated with a 1000 watts hair dryer (low wind speed mode) that produces warm air (28°–33°C) and the fluctuation of wind. The hair dryer was positioned beneath the nozzle of the wind tunnel upward to avoid direct exposure to hot wind from the hair dryer (see S4). As soon as spiders showed tiptoe behavior, the hair dryer was turned off. The turbulent intensity of the wind tunnel was 1.1% (without the hair dryer) and 11.3% (when the hair dryer turned on). There was difference in temperature between the laboratory and the field. The difference can be explained as follows: First, the ballooning behavior is coupled not only with weather condition but also with biological condition, e.g. seasonal dispersal, mating and insufficient resources, etc [2,4,26,30]. If the biological pressure, for spiders to disperse, is high, spiders may try to disperse even though it is low temperature. Second, the sudden increase of temperature acts on ballooning behavior as an influential factor [31–35]. If there is the sudden increase of temperature, e.g. because of sunshine in the morning, the ballooning behavior can be triggered even in low temperature condition. There are some reports that spiders ballooned also at relatively low temperature, 10°–20°C [33,34; the author’s field observation (13°–19°C)].

**Fig. 1.**
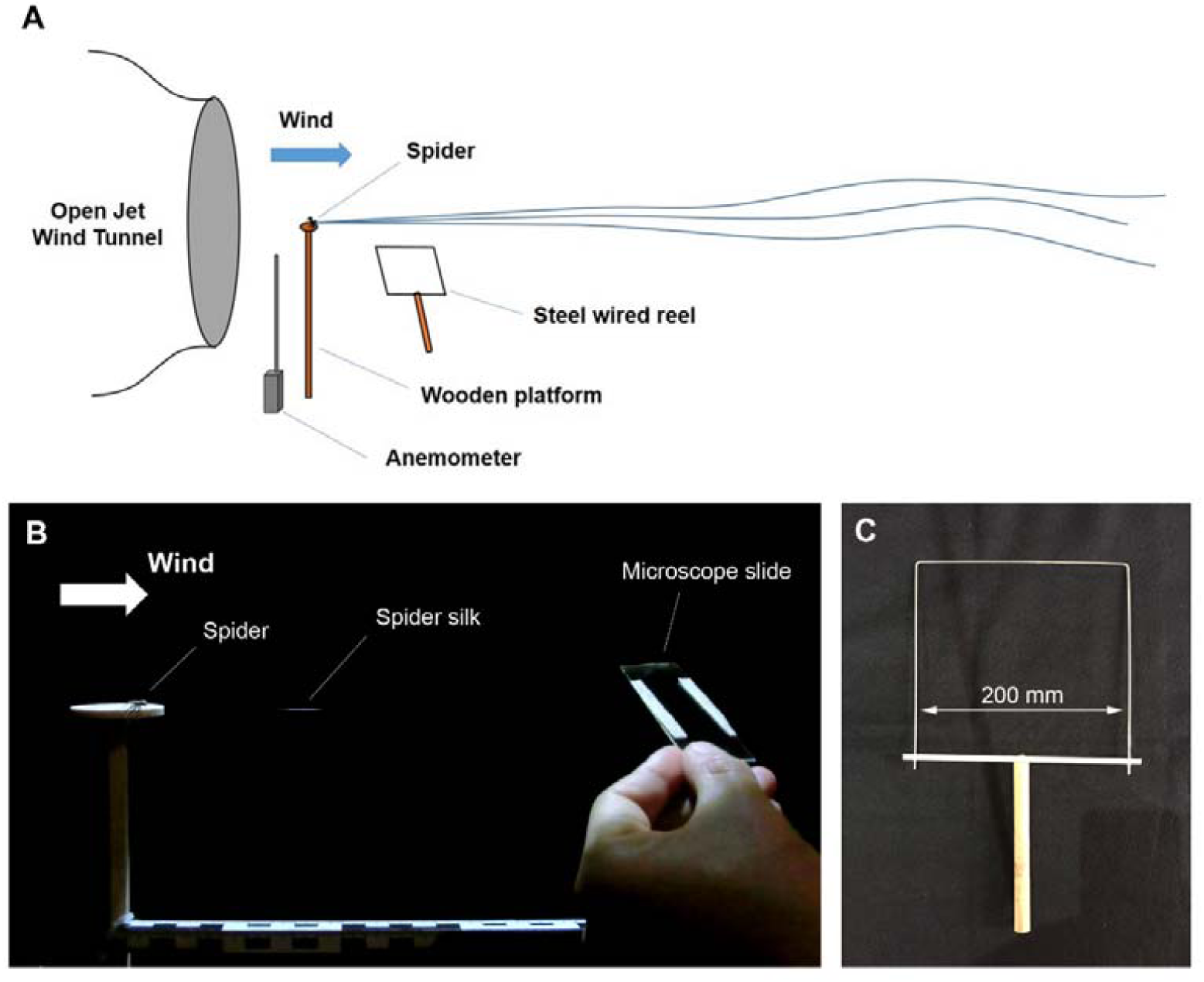
(A) A schematic view of wind tunnel tests. (B) Sampling of ballooning fibers in front of an open jet wind tunnel. (C) Reel with a steel wire to measure the length of ballooning silks.

Ballooning fibers were collected on a microscope slide, on which two narrow strips of a double-sided bonding tape were attached. The ballooning fibers were sampled near the spinnerets (see Fig. 1B). A total of 11 samples were prepared from 28 spinning events of ballooning silks. (In the experiment, a total of four spiders responded to show ballooning behavior. Two of them were very active.). Two samples were selected, because the other 9 samples failed to capture all the ballooning fibers on a single microscope slide or the fibers on the slide were deranged during the capturing process. Simultaneously, silk fibers were captured on a square wire frame and carefully wound around it in order to measure the length of ballooning threads (see Fig. 1C). The length of the silks was calculated by multiplying the total number of half revolutions by the width of the square wire frame (20 cm). The successfully sampled ballooning fibers were later studied with a field emission scanning electron microscope.

#### 2.2.2 Field emission scanning electron microscopy (FESEM)

The sampled ballooning fibers were coated with gold using a sputter coater (SCD 030, Balzers Union) and observed with a field emission scanning electron microscope (DSM 982 Gemini, ZEISS, with 5–10 kV accelerating voltage). The number of ballooning fibers was carefully counted and the thickness of fibers was measured. The spinnerets of a female *Xysticus cristatus*, were also observed with the FESEM. For sample preparation, the female spider was fixed in 2.5% glutaraldehyde and dehydrated in ascending concentrations of ethyl alcohol from 30% to l00% (10 min at each concentration). After dehydration, the sample was dried with a critical point dryer (CPD 030, BAL-TEC). The prepared sample was coated with gold using a sputter coater (SCD 030, Balzers Union) and observed with the FESEM (SU8030, Hitachi, with 20 kV accelerating voltage).

### 2.3 Investigation of the aerodynamic environment

For the investigation of the turbulent atmospheric boundary layer, ultrasonic three-dimensional wind speed measurement took place on a grass field (53° 11’ 42” N, 12° 09’ 40” E, see Fig. 2A, B). This place is also a habitat of the *Erigone* and *Xysticus* genus which do also ballooning behavior. To avoid mechanically induced updrafts by hills, trees and rocks, a flat grass field was selected. The three-dimensional wind speed data at different two mean wind speeds (1.99 m s^-1^ and 3.07 m s^-1^) were measured on a sunny day in the autumn for 5 min with the sampling frequency of 20 Hz. In order to eliminate small-scale fluctuation in the turbulent boundary layer, a quadrant analysis was introduced [40–44] (see equation (3), *u* and *w* are, respectively, horizontal and vertical wind speeds. *u’* and *w’* mean, respectively, streamwise fluctuating velocity and vertical fluctuating velocity. Those are equal to the subtracted values of the actual values of velocity minus the mean values of wind speed. *H* is the threshold parameter, which means the size of a hole. 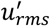 and 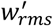 indicate root mean squared fluctuation velocities.).

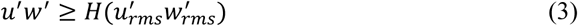

**Fig. 2.**
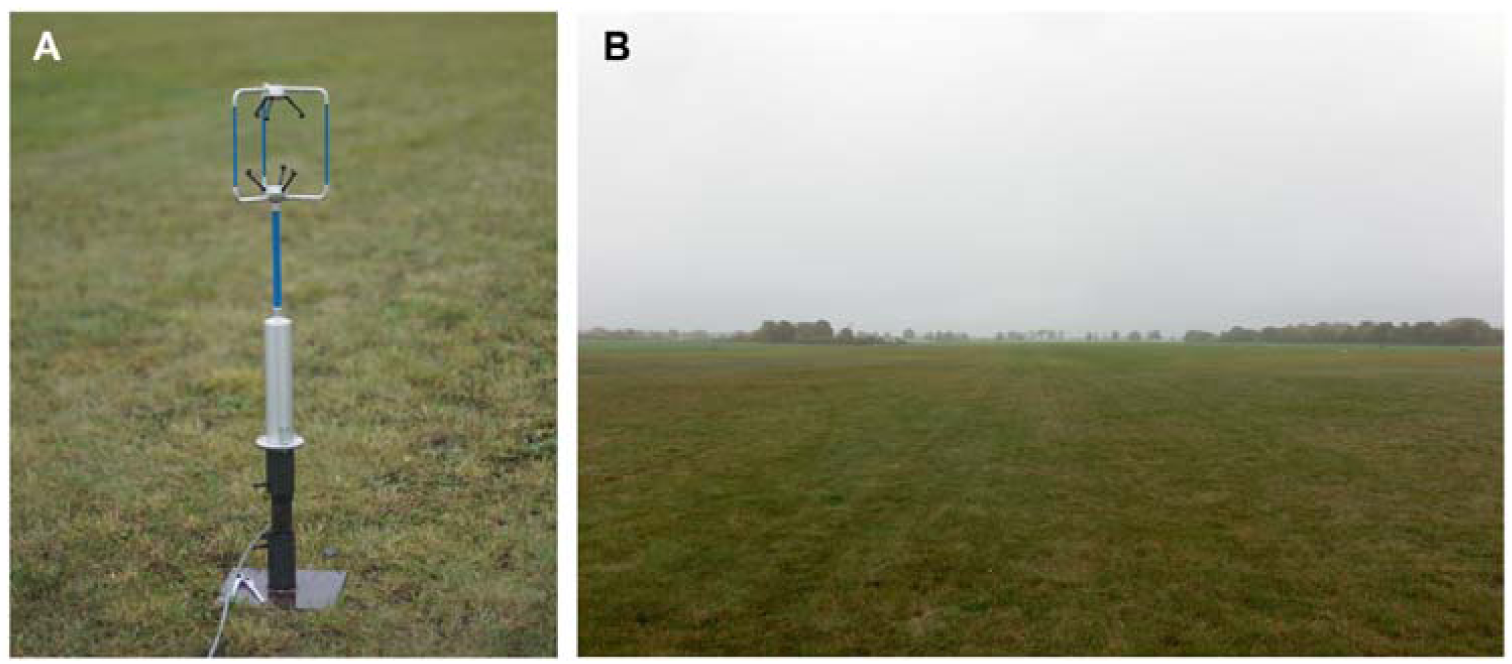
(A) A three-dimensional ultrasonic anemometer (Windmaster 1590-PK-020, Gill Instruments) is installed 0.95 m above the ground. (B) The simplest conditions (i.e. a flat surface) were selected. The flat place is covered with the 6 cm-short cut grass. Within a radius of 300 m, there is no obstacle object.

### 2.4 Animal care

Each adult and subadult crab spider (*Xysticus* genus) was raised separately in a plastic box (length × width × height: 13 × 13 × 7 cm), which has ventilation holes. Once a week, the spiders were fed a mealworm, *Tenebrio molitor*, and moisture was provided with a water spray.

### 2.5 Ethics

The species used in the experiments (*Xysticus* genus) are not endangered or protected species. No specific permissions were required. All applicable international, national and institutional guidelines for the care and use of animals were followed.

## 3. Results

### 3.1 Field observations

#### 3.1.1 Takeoff

In the *Thomisidae* family, not only female but also male adult spiders showed ballooning behaviors (see S1). During the observation days, the temperature was 16°–19°C and the mean wind speed was 6–7 m s^-1^ (gust 14–17 m s^-1^) as reported by the nearest weather station in Dahlem. The sensor was installed on 36 m position above the ground. Therefore, the local wind speed at 1.2 m above the ground must be much lower than these values. Later, we checked the wind speed on a similar day, on which the mean wind speed from the weather station showed 6–7 m s^-1^. The mean wind speed for 10 min showed 2.11 m s^-1^.

On the experiment day, the spiders mostly showed tiptoe behavior. At the first stage of tiptoe behavior, the crab spider evaluated the wind condition, not just passively through the sensory hairs on its legs, but rather actively, by raising one of its front legs (leg I) or sometimes both, and waited in this position for 5–8 sec (see Fig. 3A, B, C). This sensing behavior was often repeated a few times before the tiptoe pose. After each sensing step, the crab spider rotated its body in the direction of the wind (see Fig. 3C, D).

**Fig. 3.**
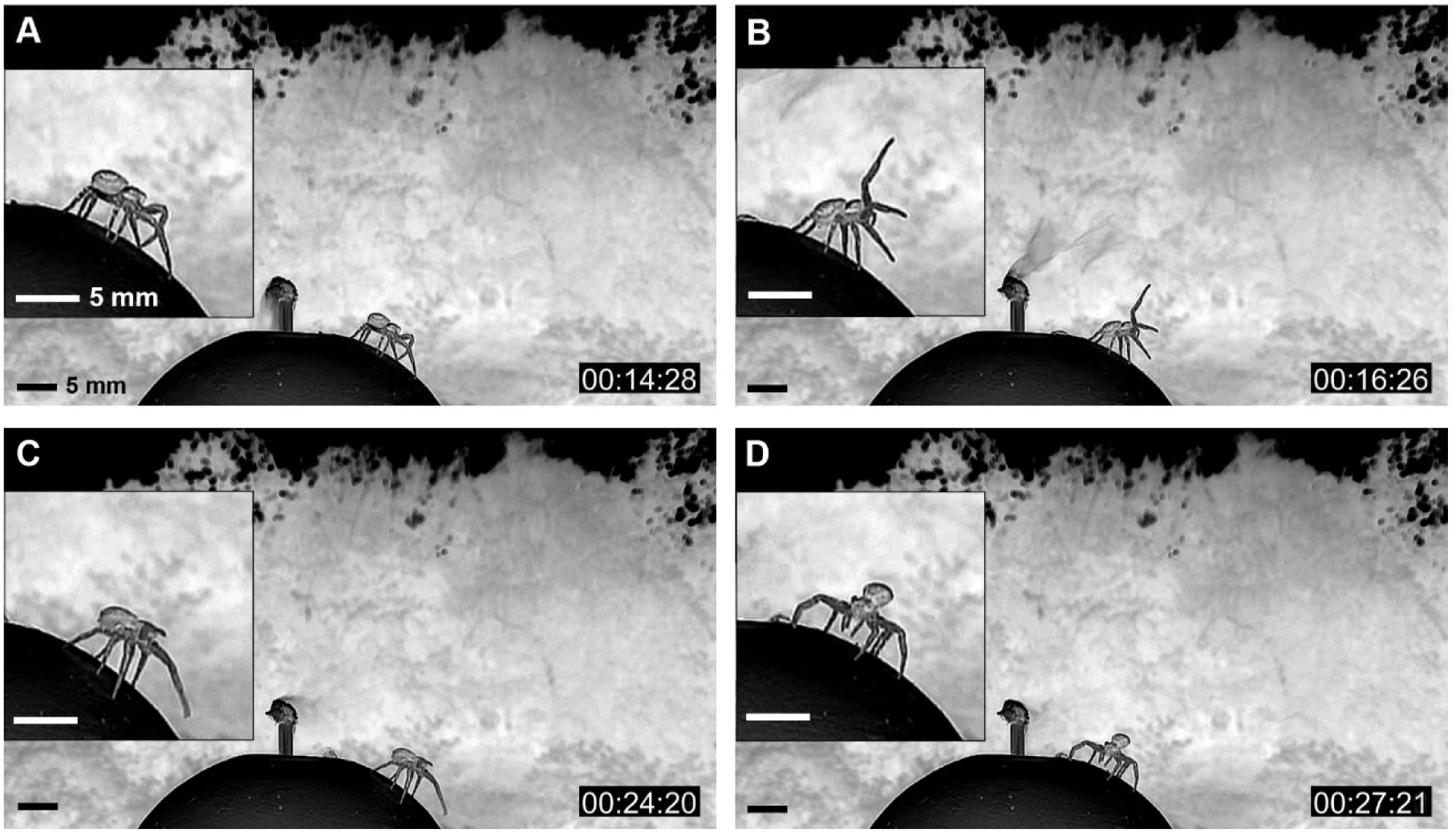
Sequence of active sensing motion with front leg (leg I) (negative images). (A) The spider first senses the condition of the wind current only through sensory hairs on its legs. (B) Then, if the condition seemed appropriate, the spider sensed more actively by raising its leg I and keeping this pose for 8 sec. (C) If the spider decided to balloon, it altered its posture. (D) The spider rotated its body in the direction of the wind and assumed tiptoe posture.

If the spider decided that the wind was adequate to balloon, it raised its abdomen (already known as a tiptoe behavior, see Fig. 3D) and spun its ballooning silks without any help from its legs. Before spinning ballooning silks, there was a motion of a rear leg (leg IV) (see S5), with which the spider holds its safety line that connected its spinnerets to the substrate, and then put it on the substrate (see S2A, B).

The crab spider first spun a single or a few number of fibers, and then many fibers (see Fig. 4A–G). The spun ballooning fibers were approximately 2–4 m long and formed a triangular sheet, which fluttered among the turbulent flows of wind. The vertex angle of this triangular sheet was about 5–35 degrees (see Fig. 4C–G). If the wind condition was not appropriate, the spider cut the silk fibers and spun them again. If the ballooning silks generated enough drag, the spider released the substrate and became airborne (see Fig. 4H). From careful video investigation, it was observed that spiders stretched their all legs outwards, as soon as the spiders achieved takeoff. Many ballooned crab spiders soared diagonally upwards along the wind flows. This paths had 5–20 degrees inclination above the horizon. Some spiders traveled quasi-horizontally. Some spiders soared along a steep path (about 45 degrees). During this steep takeoff, the spiders took off with relatively slow speed. The anchored drag line (safety line) between the platform and the spider’s spinneret could be seen. This anchored line endured without breaking, until it became 3–5 meters long. After a while, it was broken mechanically. From the wind tunnel experiment, it was found that the anchored line consists of two fibers.

**Fig. 4.**
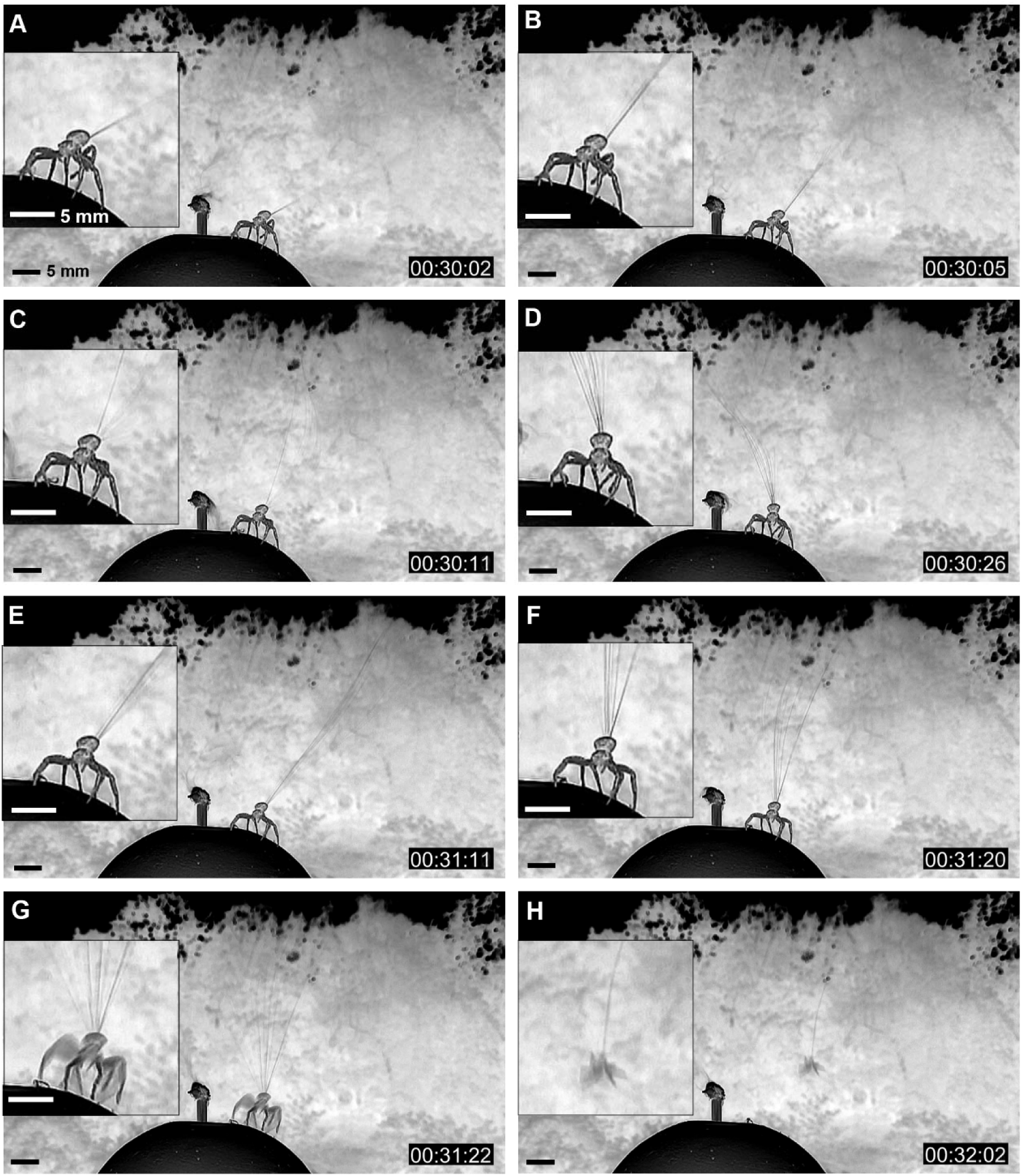
A crab spider’s ballooning process (images were converted to negative images to visualize ballooning lines). (A, B) Initial phase of spinning ballooning lines; (C, D, E, F) Fluttering of a bundle of ballooning lines. Because of turbulent flows in wind, the ballooning threads fluttered unsteadily; (G) Takeoff moment; (H) Airborne state of a ballooning spider. (Original video: see the movie S10 in supporting information.)

Three new facts about ballooning were uncovered. First, the crab spider does not evaluate the wind condition passively, but actively, by raising one of its front legs (leg I). Second, this adult ballooner anchors its drag lines on the platform not only during its rafting takeoff, but also during tiptoe takeoff. Third, the crab spider postures all its legs outwards and stretched, when airborne, not only at the takeoff moment, but also during the gliding phase (see S6A–D).

Rafting pre-ballooning behavior was also observed. The local weather condition was a little bit colder and windier than that of the previous observation day of tiptoe takeoff. Crab spiders were not active on that day. As soon as they were set on the platform, they showed one of two behaviors. Either they hid on the opposite side of the platform to avoid the wind, or they quickly retreated downwards about 0.4 to 1.1 m relying on their drag lines (see S3A, B), and spun their ballooning fibers downstream of the wind. At this time, spiders spun a single or a few number of fibers first, and then many fibers, as they showed during the tiptoe takeoff (see S3C, D). During this process, the spiders also postured all their front legs and second legs outwards and backwards, so that they hung and directed their bellies in an upwards direction of the wind. The backward (downstream) spun threads slowly curved upwards and the spiders’ bodies also slowly moved upwards (see S3D). At some point, when the threads had generated enough drag and lift, the drag lines near the spinnerets were cut and the spider finally ballooned (see S3E, F).

#### 3.1.2 Statistical analysis of pre-ballooning behaviors

The spiders showed 67 active sensing motions with their leg I. Forty-four tiptoe pre-ballooning motions could be observed. Six spiders had ballooned successfully after their tiptoe pose (see S7). They also dropped down 8 times relying on their drag line. Three of them showed a rafting takeoff (see S7).

The initial behaviors were mostly started with sensing motion. The frequent transitions between different behaviors were occurred between sensing motion and tiptoe motion (see Fig. 5A). Spiders flew from only either the tiptoe pose or the dropping and hanging pose (see Fig. 5B). The probability of the ballooning takeoff from tiptoe behavior was 9.5%. The probability of the ballooning takeoff from the rafting pose was 37.5%.

**Fig. 5.**
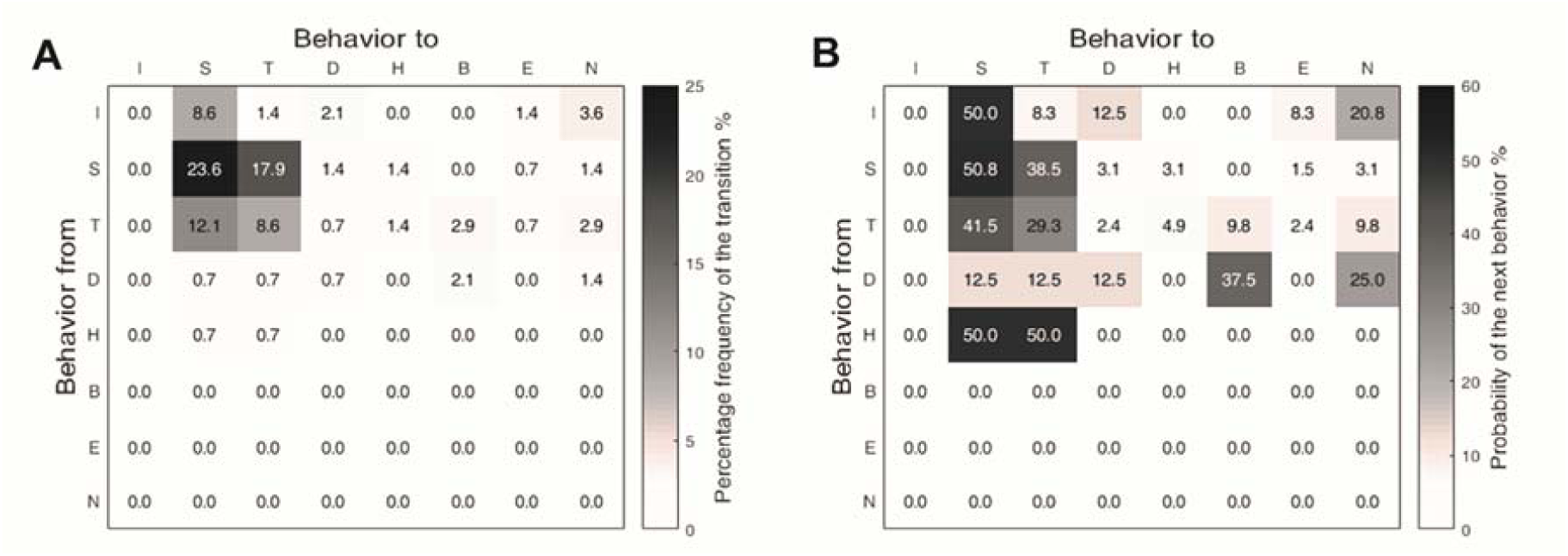
(A) The percentage frequency of the behavior transition (the total number of transitions: N = 144; I: initial state, S: Sensing motion, T: Tiptoe behavior, D: Dropping and hanging behavior, H: Hiding motion, B: Takeoff, E: Escape, N: Not flown). (B) The transition matrix between behaviors (the total numbers of categorized behaviors: *N*_*I*_ = 25, *N*_*S*_ = 67, *N*_*T*_ = 42, *N*_*D*_ = 8, *N*_*H*_ = 2).

The duration of each tiptoe behavior was measured and their frequencies were analyzed. Short period tiptoe poses, which lasted for less than 5 sec, were the most frequent. The longest tiptoe event lasted 65 sec. Successful ballooning takeoffs were not biased in relation to tiptoe duration (see Fig. 6).

**Fig. 6.**
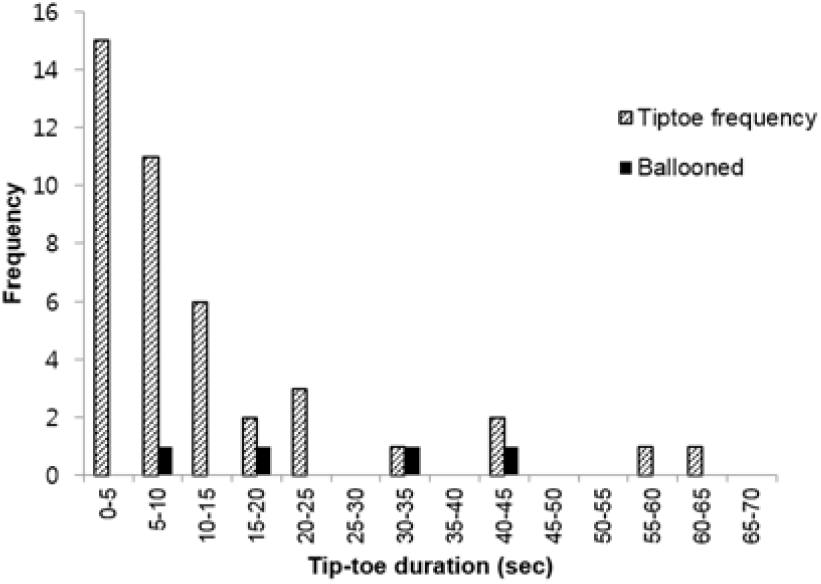
Frequency diagram of tiptoe behaviors according to tiptoe duration (N = 42). Black columns are the tiptoe behaviors that were connected to the successful ballooning takeoffs (N = 4).

#### 3.1.3 Gliding

A total of 32 floating threads were observed at the Teltow canal. Most of them were horizontally transported along the channel at about 1–8 m above the water surface. They drifted passively due to light wind, but rarely fell down. Some of their silks were inclined downstream. The others were inclined upstream. Two of 32 floating threads were just threads alone without a spider. The number of observed threads was one to five. However, as not all threads were visible with the naked eye, some may have been the multiple threads, which stuck together, although they seemed to be a single thread. Some of them may not have been seen, because of their inappropriate angles and positions in relation to the sun. Most of the spiders were positioned at the lower end of their threads. Although the threads showed different numbers and shapes, they were usually laid diagonally (see Fig. 7).

**Fig. 7.**
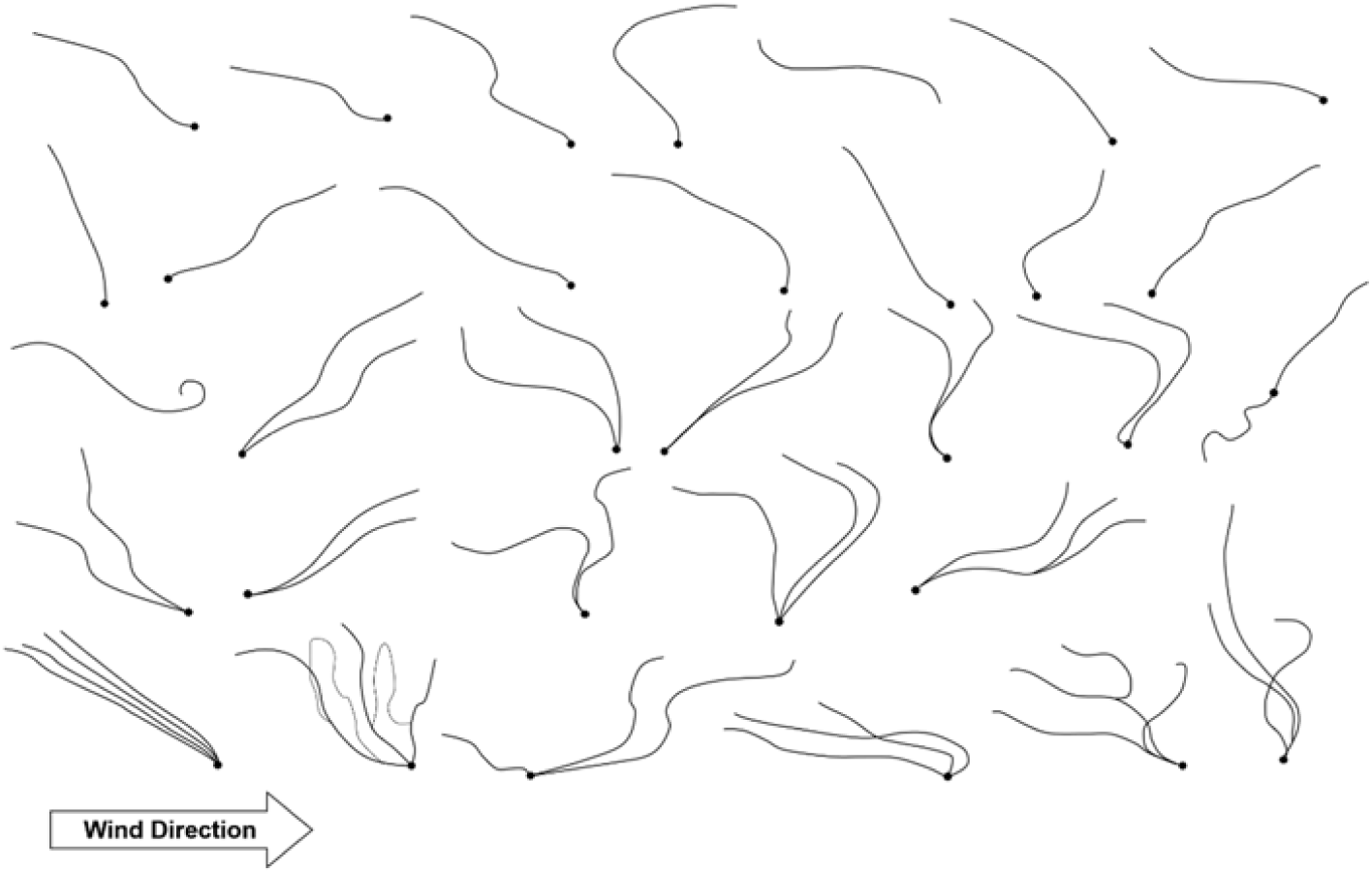
Sketches of ballooning structures (body + ballooning threads). These structures were observed above the water surface, at heights of 1–8 m. Wind was blowing from left to right. Therefore, these structures were transported in the same direction as the wind. Black and thick points represent spider’s body. Black lines represent ballooning threads.

### 3.2 Identification of ballooning fibers

The separation of glands for a drag line and ballooning lines was observed in ballooning behaviors in the laboratory. The anchored drag line was connected to the anterior spinnerets and ballooning fibers were spun from either one or both posterior or/and median spinnerets (see S5). The lengths of the ballooning fibers were measured, 3.22 ± 1.31 m (N = 22), from 22 spinning events of two crab spiders (16.4 and 18 mg) (see Fig. 8). The maximum length of the spun ballooning lines was 6.2 m.

**Fig. 8.**
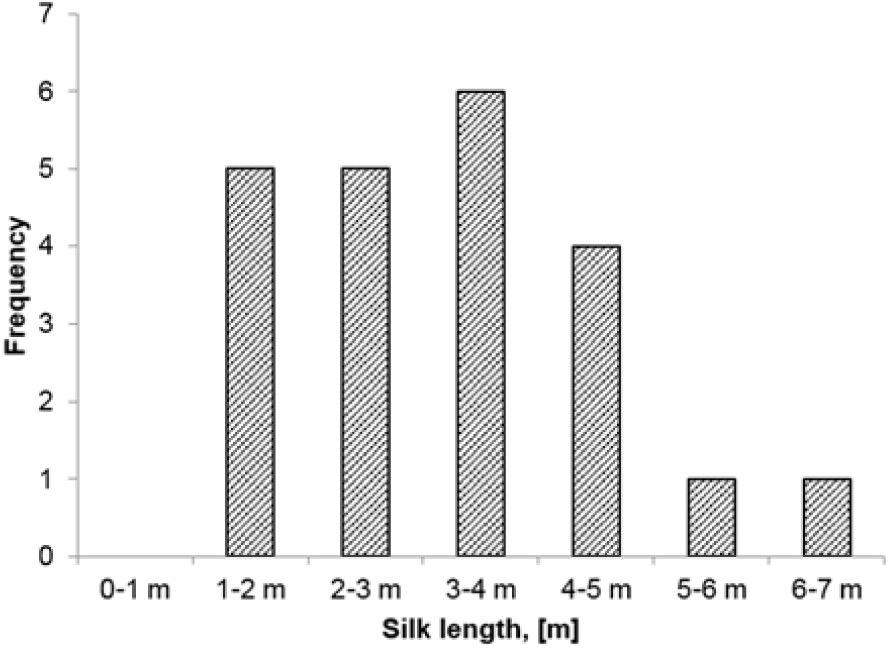
Distribution of the length of ballooning lines (N = 22).

The successfully collected ballooning fibers of both *Xysticus cristatus* and *Xysticus* species were observed with the FESEM. Ballooning fibers consisted of two thick nanoscale fibers that were attached together (see Fig. 9B, C) and many thin nanoscale fibers (see Fig. 9A–D). The two adult spiders, *Xysticus* species, spun, 48 to 58 thin nano-fibers and 2 thick nano-fibers (see Table 1). The thickness of the thin nano-fibers ranged from 121 to 323 nm with an average of 211.7 ± 45.2 nm (N = 40). The thickness of the thick nano-fibers was 698 to 768 nm with an average of 722.2 ± 32.5 nm (N = 4) (see Fig. 9B–D). The spun fibers were split independently (see Fig. 9A–D).

**Table 1.** Identification of the number and thickness of ballooning fibers through FESEM.

| Species | Weight | Thin nanoscale fibers |  | Thick nanoscale fibers |  |
| --- | --- | --- | --- | --- | --- |
|  |  | Number of fibers | Thickness of fibers | Number of fibers | Thickness of fibers |
| **Spider 1** *Xysticus spec. (f.)* | 20.8 mg | 58 | $192.3 \pm 36.3$ nm (N = 20) | 2 | $734.0 \pm 48.1$ nm (N = 2) |
| **Spider 2** *Xysticus cristatus (f.)* | 18 mg | 48 | $231.1 \pm 45.6$ nm (N = 20) | 2 | $710.5 \pm 17.7$ nm (N = 2) |
| **Average** | - | $53 \pm 7.1$ (N = 2) | $211.7 \pm 45.2$ nm (N = 40) | 2 (N = 2) | $722.2 \pm 32.5$ nm (N = 4) |

**Fig. 9.**
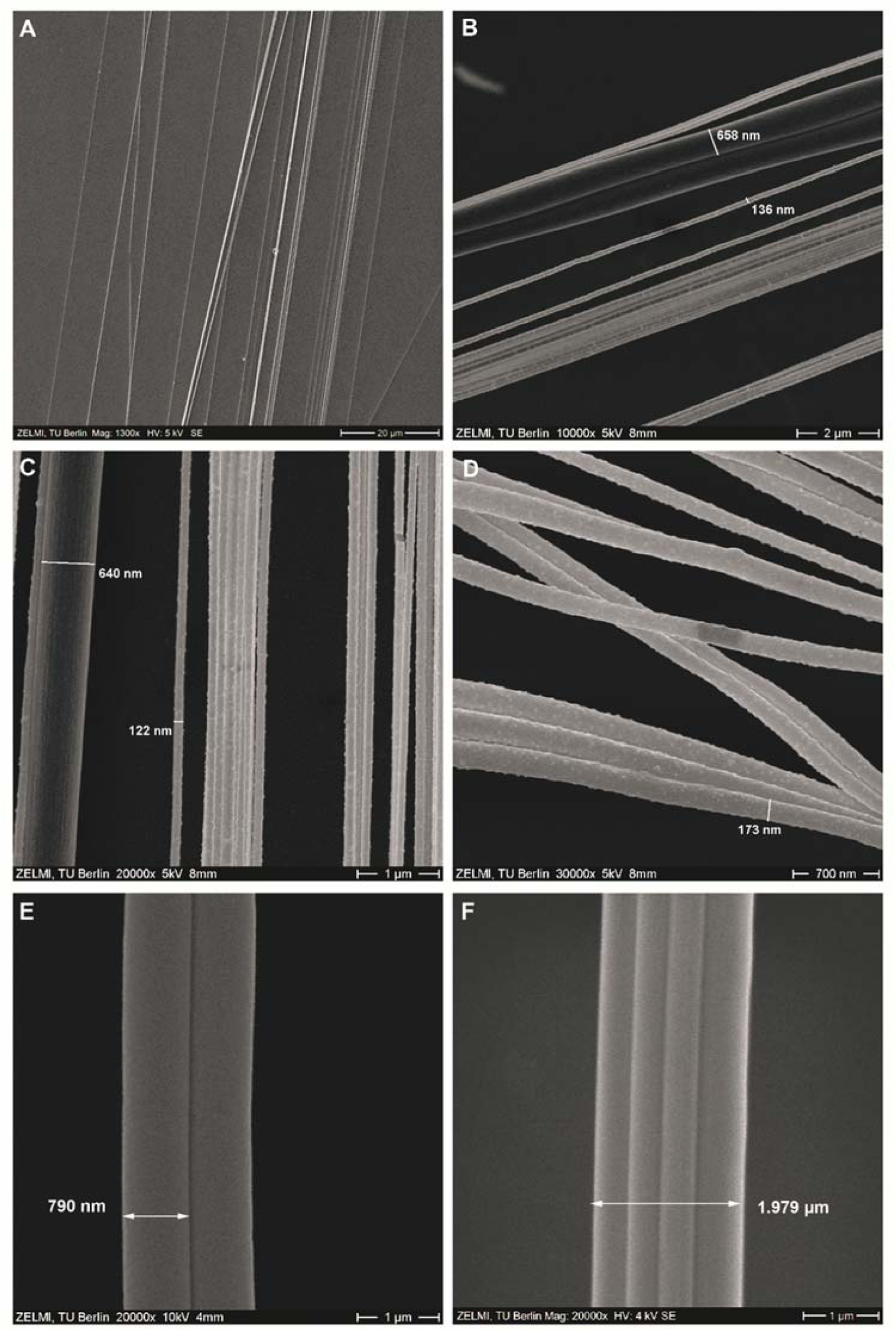
(A) Ballooning fibers of *Xysticus cristatus*. (1300×). (B) Ballooning fibers of *Xysticus audax* (10000×). (C) Middle part of ballooning fibers of *Xysticus audax* (20000×). (D) A bundle of ballooning fibers of *Xysticus audax* at the end of threads (20000×). (E) One pair of drag fibers of *Xysticus cristatus* (a weight of 18 mg) (20000×). (F) Two pairs of drag fibers of *Xysticus* species (a weight of 15.6 mg), which attached together (20000×).

On the other hand, drag lines consist of one pair of fibers (see Fig. 9E) (sometimes two pairs, from left and right, see Fig. 9F) which were spun from the major ampullate glands on the anterior spinnerets. The thickness of these major ampullate silks was about 790 nm for *Xysticus cristatus* and 493 nm for *Xysticus audax*, respectively (see Fig. 9E, F).

### 3.2 Aerodynamic environment on the short grass field

Two data sets at the different mean wind speeds (1.99 m s^-1^ and 3.07 m s^-1^) for 5 min were collected. Usable updrafts were investigated in the turbulent atmospheric boundary layer. Each of the cases showed the vertical deviation of ±0.225 m s^-1^ and ±0.267 m s^-1^, respectively, which ranged from −0.5 to 0.5 m s^-1^ and from −0.6 to 0.7 m s^-1^ (see Fig. 10A, B). The turbulent intensities were 21 % and 23.7 % in the mean wind speeds of 1.99 m s^-1^ and 3.07 m s^-1^, respectively. The vertical wind speed of both cases is negatively correlated to the horizontal wind speed. The slope of regression fits are −0.235 (r = −0.575) and −0.077 (r = −0.259), respectively. The horizontal wind speeds of both Q2s are positioned below the wind speed of 3 m s^-1^.

**Fig. 10.**
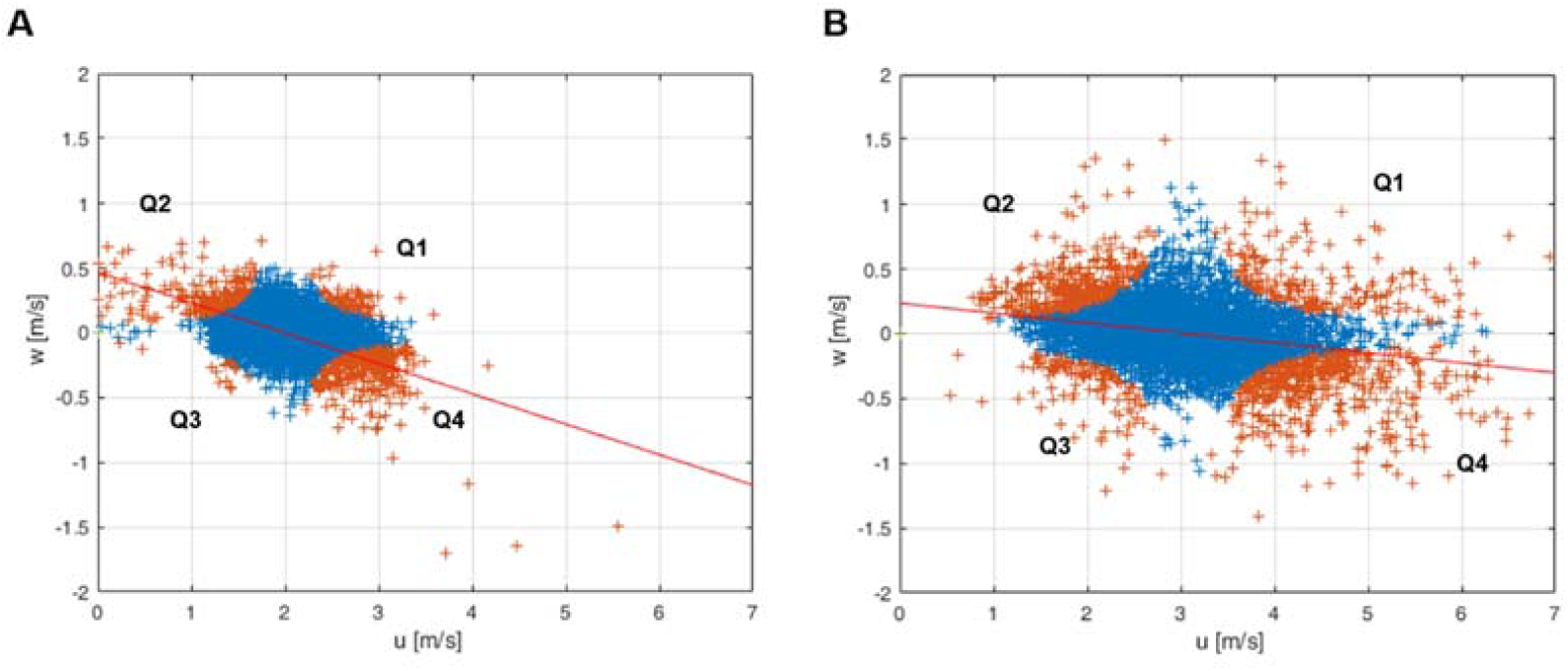
Horizontal and vertical components of the wind speeds for 5 min on *u – w* domain. (20 Hz sensing rate). Orange cross points: The quadrant data (Q1–Q4) of the measured wind speeds according to *u’* and *w’*. Blue cross points: The ignored data regarding as a small-scale fluctuation (*H <* 1). Red lines are linear regression fit lines. (A) The case of 1.99 m s^-1^ mean wind speed. (30th October 2016 12:39 – 12:45 LT) (B) The case of 3.07 m s^-1^ mean wind speed. (29th October 2016 10:54 – 11:00 LT)

## 4. Discussion

From our observation, the physical properties of ballooning silks were identified together with previously undescribed behaviors during ballooning: (i) an active sensing motion of the wind, (ii) an anchoring behavior during tiptoe takeoff, (iii) a tidying-up motion of an anchor line (drag line, safety line) and (iv) an outward stretching pose during a flight. These findings provide some clues and answers, which are previously unsolved in spiders’ ballooning flight. The measured wind data also provide the possible mechanism from the viewpoint of the environment. We interpret as follows.

### Spider’s active sensing motion of the wind condition

**Crab spiders showed the motion that seemed like actively evaluating the meteorological condition before their takeoff.** Normally ballooning behavior is first triggered either by a warm ambient temperature or by a rapid increase in ambient temperature [31,36]. Additionally, if they are exposed to a wind that is slower than 3 m s^-1^ (favorable in 0.35–1.7 m s^-1^ wind speed for spiderlings), they show tiptoe behavior [1,3,36,37]. Until now, it had been thought that spiders sense the wind speed passively through the sensory hair (Trichobothria) on their legs [17,31,45]. However, the present observations show **that spiders may sense the aerodynamic condition of wind not just passively, but rather actively, raising their leg I high and shaking it.** Strong interactive behavior transition between sensing motion and tiptoe motion (see Fig. 5A) supports that active sensing motions are related to the spiders’ ballooning flight, because the tiptoe behavior is prerequisite behavior for ballooning takeoff (see Fig. 11). The leg-raising behavior can be interpreted as follows: the spider enhances the sensibility of its sensory hairs, by raising its legs upward in the outer region of the boundary layer, where airflows are faster than near the substrate and undisturbed by spiders’ body themselves. The first instars of the *coccid Pulvinariella mesembryanthemi* show a similar behavior by standing on their hind legs to capitalize on higher air speed in the thin boundary layer for their aerial dispersal [46].

**Fig. 11.**
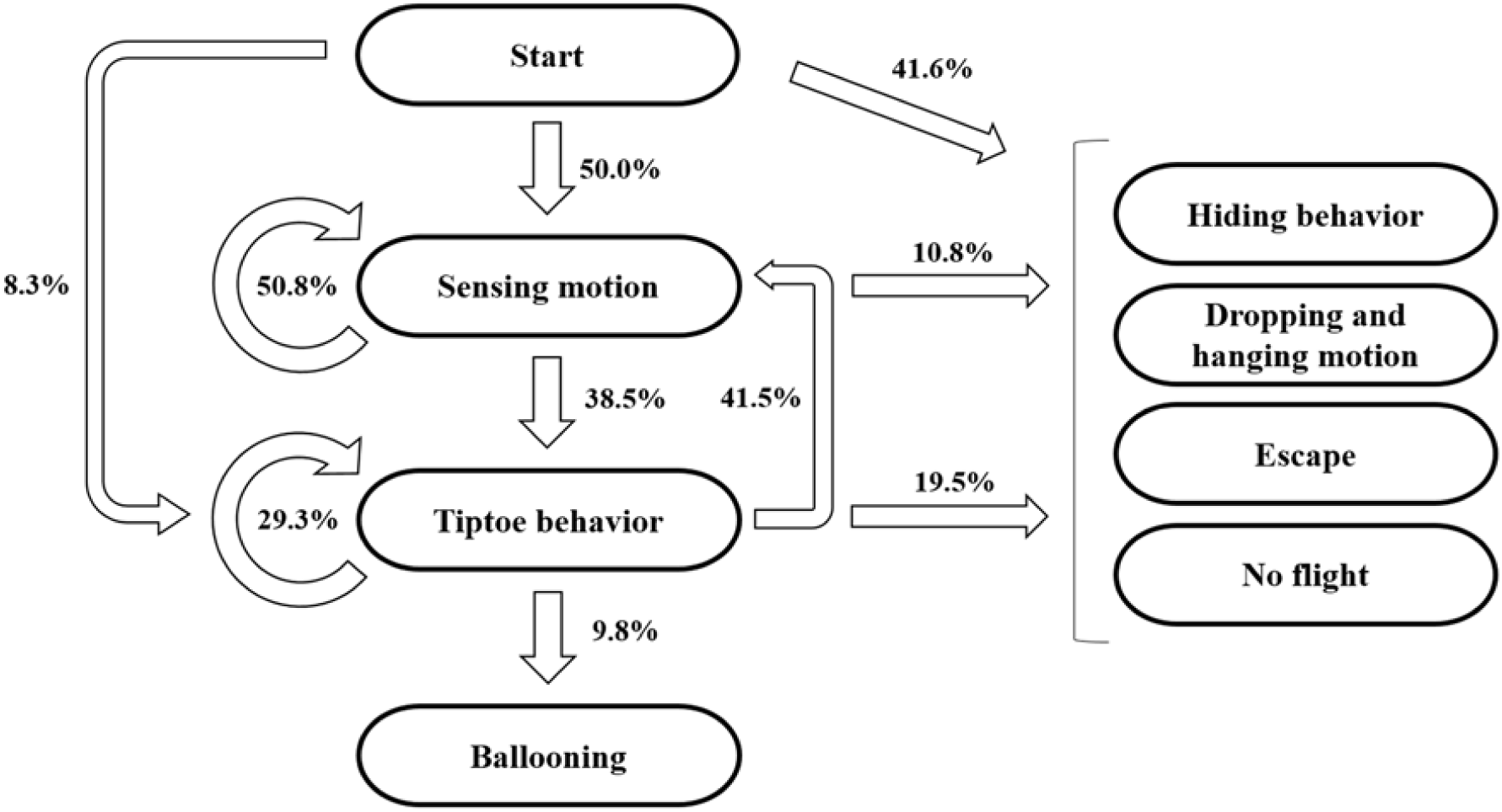
Takeoff process in tiptoe ballooning. The probabilities are calculated based on the total number of behaviors at each stage (see Fig. 5B).

The spider’s active sensing motion of the wind tells us two important things. **First, in spiders ballooning, aerodynamic force maybe be a dominant factor.** One hypothesis claims that an electrostatic charge on ballooning silks could generate lifting forces in the Earth’s vertical atmospheric electrostatic field [22]. However, the leg-raising behavior indicates that, from the spider’s viewpoint, airflow is an important factor for its ballooning. **Second, the spiders’ ballooning may be not random flight that simply relies on the random condition of the wind, but they may sense and evaluate the condition of the wind and wait for the appropriate moment to initiate ballooning.** Thirty-eight percent of tiptoe behaviors lasted 10–65 sec. This can be interpreted that there may be favorable wind conditions for spiders to balloon. If spiders evaluate wind conditions and wait for their aerial dispersal, this can be a distinct feature of spiders’ ballooning in contrast to other passive aerial dispersal, like that of seeds or aero-planktons. Additionally, this evaluating behavior can also save their silk dopes during their takeoff trials by reducing failure cases, for example, by avoiding unfavorable wind conditions after spinning the ballooning silks.

In the near surface atmospheric boundary layer, the updraft is not initiated by thermal buoyancy forces but is initiated by the shear winds, which generate shear-driven turbulent flows [47,48]. The generated turbulent eddies build intermittent updraft and downdraft regions, which drift to the downwind direction. If spiders sense either this moving updraft zone directly or the appropriate wind condition, which can build this moving updraft structure, indirectly, the spiders’ sensing motion and timing the decision may be helpful for their successful takeoff.

There are still other questions that can be asked. If spiders sense these wind conditions, what type of information do spiders need for their decision to balloon? From previous studies and the author’s observations, we deduce several factors. (i) Wind speed: A spider does not show tiptoe behavior under conditions of high wind speed, over 3 m s^-1^ [1,3,36,37]. (ii) A vertical wind speed: Favorable condition for ballooning, usually vertical acceleration of wind, persisting only for a few seconds. Under such a condition, the spider spins its silk fibers in wind rapidly and releases its substrate [20,36]. (iii) Wind direction: From our observation, spiders rotate their body in the direction of the wind, as soon as they have evaluated the wind condition. Therefore, at least, spiders perceive wind direction. (iv) Wind fluctuation: For drop and swing dispersal, spiders were particularly more active with turbulent flows [49]. This shows that spiders can perceive the fluctuation of a turbulent flow. Suter showed that the ballooning site is usually laid within chaotic air flows [20]. Reynolds showed that the chaotic motion of turbulent flow reduces the terminal speed of ballooners and that this feature enables long permanence in the air [16]. Therefore, it can be deduced that spiders may sense the fluctuation of wind.

### Ballooning lines and anchored line

For crab spiders, ballooning lines were not identical with a drag line. The source of ballooning lines are not well known [3,19]. There is a report that some of primitive spiders, *Sphodros* spiderlings and *Ummidia* spiderlings, which do not use the complex pre-ballooning behaviors such as tiptoe and rafting, use a drag line as a ballooning line, which is known as “suspended ballooning” [10]. Spiders drop down from the end of a branch relying on their drag line. If there is a breeze, the drag line near its point of attachment to the platform would be sheared and drift through the air [28,50]. From this context, many previous studies regarded that spiders use their drag line for ballooning dispersal [16,17,22,23,38]. Some experiments substituted a drag line for a ballooning line [18,19,21]. However, Suter guessed from Tolbert’s observation that ballooning lines might be different from the drag line [3,18,20]. Our observation assures that in crab spiders, these ballooning lines are not identical with a drag line, because those were spun from the either or both of the median or/and posterior spinnerets (see S8A, B). Normally, drag line is spun from major ampullate glands in anterior spinneret [51]. The crab spiders spun 48 to 58 thin nano-fibers and one pair of 2 thick nano-fibers. These thin nanoscale fibers seemed to be aciniform fibers (wrapping silk) from the median/poster spinnerets. The thick nanoscale fibers seemed to be minor ampullate silks from the median spinnerets. Because Moon and An observed that one of median spinnerets (left and right) of Misumenops tricuspidatus (crab spider) contains 2 pairs of ampullate glands and 20 (±3) pairs of aciniform glands and one of posterior spinnerets of it includes 50 (±5) pairs of aciniform glands [52]. Our observation in Xysticus species showed similar features with Moon and An’s observation (see S8C, D).

The anchored drag lines are possibly normal drag lines (major ampullate silks), which a spider spins constantly while it is crawling and attaching its silk fibers to the ground surface for safety purposes. Because the anchored line was connected to the anterior spinnerets (see S8A) and was consisted of two fibres. From Osaki’s observation, drag line are also composed with two fibers [53]. The anchored line can be observed in every ballooning behavior, not only rafting takeoff [10] but also tiptoe takeoff.

Before the spinning motion of ballooning lines, there was a motion of a rear leg (leg IV) (see S5), resembling “wrap spinning”, which spiders use leg IV to initiate ballooning lines by wiping their spinnerets [53]. However, in our case, it was obvious that the spiders did use their leg IV not to initiate ballooning lines, but to tidy up the unfamiliarly positioned anchored line (safety line), which might have obstructed the spinning of ballooning silks (see S2A, B and S5A, B). During the tiptoe takeoff, this anchored line is normally cut in a short time mechanically, when a spider drifts fast either horizontally or at a small inclination. But when a spider becomes slowly airborne with a very steep inclination because of a local updraft, this line bears until it is lengthened to 3–5 m. In our opinion, it is not stretched, but a spider lengthened the safety silk by spinning an additional safety line. The reason is that the maximum elongation of spider silks is limited by about 30% [55,56]. This means that spiders control their anchored line during the steep or slow takeoff.

### Nanoscale multi-fibers and ballooning flight

While the ballooning of small spiders, which are lighter than 2.0 mg, was investigated by calculation [17], experiments [18,21] and observation [1,3], the plausibility of a large spider’s ballooning was not yet explained [17,39]. **The mysterious flying behavior of large spiders can be explained by their nanoscale multi-fibers.** From our wind tunnel test, we found that the *Xysticus* genus uses tens of nano fibers (diameters of 121 to 323nm) for their aerial dispersal (see Fig. 9A–D). The number of ballooning fibers and their lengths were identified. Based on these measured values, the required updraft speed for the ballooning takeoff was calculated using modified Humphrey’s and Suter’s equations (equation (4)–(8)) [17,18]. The fluid-dynamic interaction between each fibers is not considered. The split case of fibers are calculated. **For a crab spider weighing 10 to 25 mg, the required vertical wind velocities, according to Humphrey’s theoretical formula 0.08–0.20 m s**^**-1**^, **and according to Suter’s empirical formula, are 0.04–0.09 m s**^**-1**^. These values are much smaller than the values, 9.2–21.6 m s^-1^, which were calculated for *Stegodyphus* species [25,39]. We calculated the required length of ballooning fibers for *Stegodyphus* species according to the updraft air speeds (see Fig. 12). **Our result shows that *Stegodyphus* species can also balloon with relatively light updraft, 0.2–0.35 m s**^**-1**^, **with 3 m length and a total 80 numbering ballooning fibers.** These results support Schneider’s observation that adult females of the *Stegodyphus* genus, 80– 150mg, balloon with at least tens to hundreds of threads [26]. In this calculation, the number of ballooning fibers for *Stegodyphus* genus are assumed conservatively that only one side (left or right) of the median and posterior spinnerets are used for spinning ballooning fibers. This number of fibers is estimated by counting the fiber glands of *Stegodyphus mimosarum*’s spinnerets (the number of minor ampullate fibers is 2, the number of aciniform fibers is 78) [57]. This number is reasonable, because *Xysticus* species spun similar order of the number of fibers (2 number of thick fibers and 58 number of thin fibers from our experiments). The thicknesses of a minor ampullate fiber and an aciniform fiber are assumed 80% and 25% of that of the major ampullate fiber, respectively. The thickness of a major ampullate fiber is estimated from Suter’s body weight and silk thickness relationship (equation (8), Suter 1991).

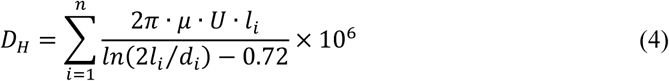

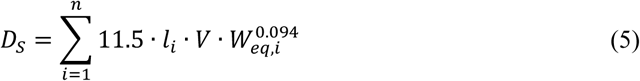

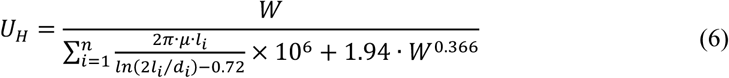

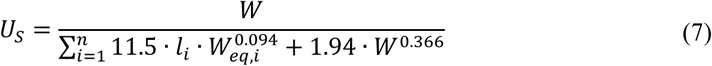

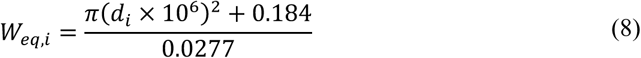

*D*_*H*_: Drag of multiple fibers by Humphrey’s equation in *µN*.

*D*_*S*_: Drag of multiple fibers by Suter’s equation in *µN*.

*U*_*H*_: Required vertical wind speed for ballooning using Humphrey’s equation in m s^-1^.

(The spider’s body drag is used from the Suter’s empirical relationship.)

*U*_*S*_: Required vertical wind speed for ballooning using Suter’s equation in m s^-1^.

*µ*: Dynamic viscosity of the air at 20°C (1.837 × 10^−5^*kg · m*^−1^ *· s*^−1^).

U: Velocity of the air in m s^-1^.

*l*_*i*_: Length of i-th silk fiber in m.

*d*_*i*_: Diameter of i-th silk fiber in m.

*W*: Weight of the spider body in *µN*.

*W*_*eq,i*_: Equivalent weight of the spider body corresponding to the diameter of a ballooning fiber in *µN*.

**Fig. 12.**
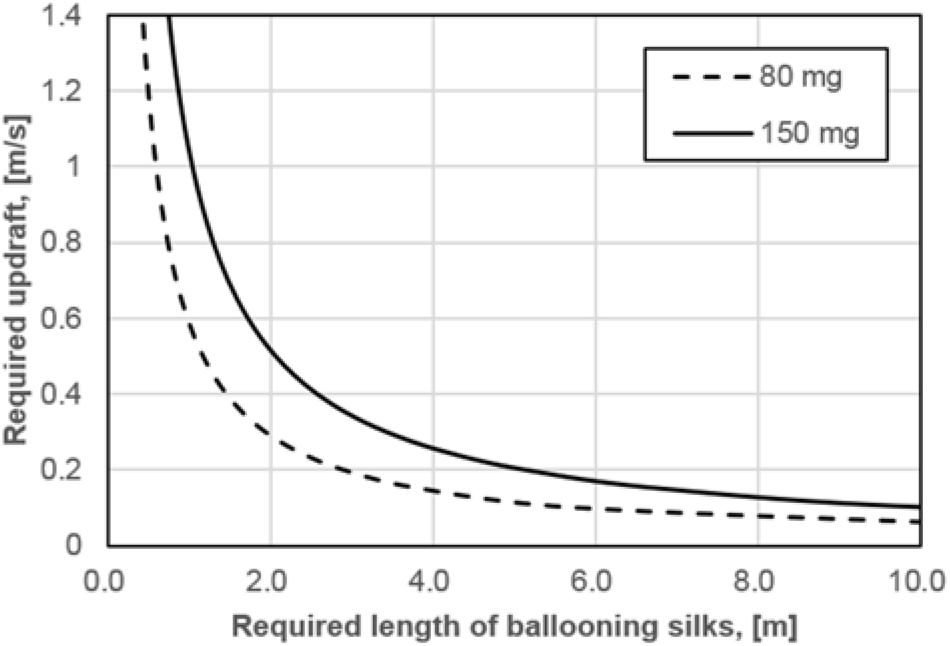
Required updraft wind speed and length of ballooning silks for the ballooning of 80–150 mg *Stegodyphus* species. It is assumed that *Stegodyphus* species use 2 minor ampullate silks (2.1–2.9 µm thickness) and 78 aciniform silks (650–900 nm thickness) for their ballooning.

### Low Reynolds number fluid-dynamics in a spider’s ballooning flight

The ballooning silks, exposed to the air, experience a low Reynolds number flow (Stokes flow, their Reynolds number is lower than 1). The upper limit of the Reynolds number is about 0.04 (*Re = ρVd/µ*; air density: 1.225 kg m^-3^, maximum possible velocity: 3 m s^-1^, thickness of spider silk: 211 nm, dynamic viscosity of air: 1.837 × 10^−5^ kg m^-1^ s^-1^). The maximum possible speed that the thread experiences is assumed to be 3 m s^-1^, because a spider seldom flies above this wind speed. Once a spider is airborne, the relative air speed with respect to the silk is reduced. Therefore, the Reynolds number of spiders’ silks during their flight is much smaller than 0.04, and the spider’s flight is dominated by low Reynolds number fluid-dynamics [58]. However, a large spider’s body shows much larger scales than those of a spider silk, not only in size, but also in weight. In a free-fall case of a spider body without any silks, the Reynolds number is about 2300 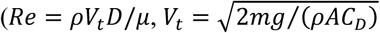; a spider’s body is assumed as a 5 mm diameter sphere with 25 mg weight, the projected area of a sphere: 19.6 *mm*^2^, the coefficient of a sphere: 0.43, the terminal speed: 6.2 m s^-1^) [59]. In this Reynolds number flow regime, **the drag of a body is proportional to almost the square of relative wind speed**, because the Reynolds number region of 5 *< Re <* 3000 is categorized as moderate Reynolds numbers, which inertial force of flow is still dominant in comparison with a viscous flow [60]. On the other hand, **the drag of spider silk is proportional to relative wind speed**, because a viscous force is dominant in this low Reynolds number regime (*Re ≪* 5) [60]. **Therefore, the macro-scale spider’s body, which means that a spider falls with acceleration without ballooning silks, is suspended by their high tensile strength nanoscale fibers that experience micro-scale fluid-dynamics (low Reynolds number flow). This enables a spider to float in the air like a particle which experiences relatively low Reynolds number flow.** The Reynolds number of a spider’s body with ballooning fibres is about 30 (*Re = ρVD/µ*; minimum possible velocity: 0.09 m s^-1^ by the lowest terminal speed from modified Suter’s formula). The utilization of low Reynolds fluid-dynamics in ballooning flight is different from flight mechanics of other winged insects’ flight, which mostly uses moderate or high Reynolds number aerodynamics, 10^3^ *< Re <* 10^5^ [61-63].

### Low wind speed and turbulence

**The measured vertical wind speeds of updrafts ranged 0–0.5 m s**^**-1**^ **at the mean wind speed of 1.99 m s**^**-1**^ **and 0–0.7 m s**^**-1**^ **at the mean wind speed of 3.07 m s**^**-1**^**. These values are enough for spiders’ ballooning, the required vertical wind speeds of which were 0.04–0.2 m s**^**-1**^ **for 10–25 mg *Xysticus* species and 0.2–0.35 m s**^**-**^**1 for 80– 150 mg *Stegodyphus* species.** The observed negative correlation between vertical wind speed and horizontal wind speed in the turbulent boundary layer, which has been also studied in the fields of fluid-dynamics and meteorology [42–44], suggests to us that **the reason spiders show their pre-ballooning behavior at a low wind speed regime (smaller than 3 m s**^**-1**^**) is because they may use the “ejection” regimes in turbulent flow, which contain updraft components and are induced by a “coherent structure” near the surface boundary layer.** A shear wind on a planted field, a meadow or relatively short plants, forms an organized structure called the “coherent structure” (possibly “hairpin vortex” or “horseshoe vortex,” see Fig. 13A, B; “hairpin vortex packets” over relatively short plants and “dual hairpin vortex structures” over planted field (see Fig. 13C, D)) [42,64–68]. These structures intermittently produce up- and downdrafts [42,65,67]. The interesting point is that these updrafts are highly correlated with a decrease in wind speed, which is categorized as Q2 (*u’ <* 0 and *v’ >* 0), an “ejection” region in a quadrant analysis [42–44] (see Fig. 10). Therefore, the phenomenon that spiders usually balloon in the low wind speed regime (lower than 3 m s^-1^) could be explained with this organized structure in the atmospheric turbulent flows above the ground. However, the frequency and duration of these updrafts are not well known.

**Fig. 13.**
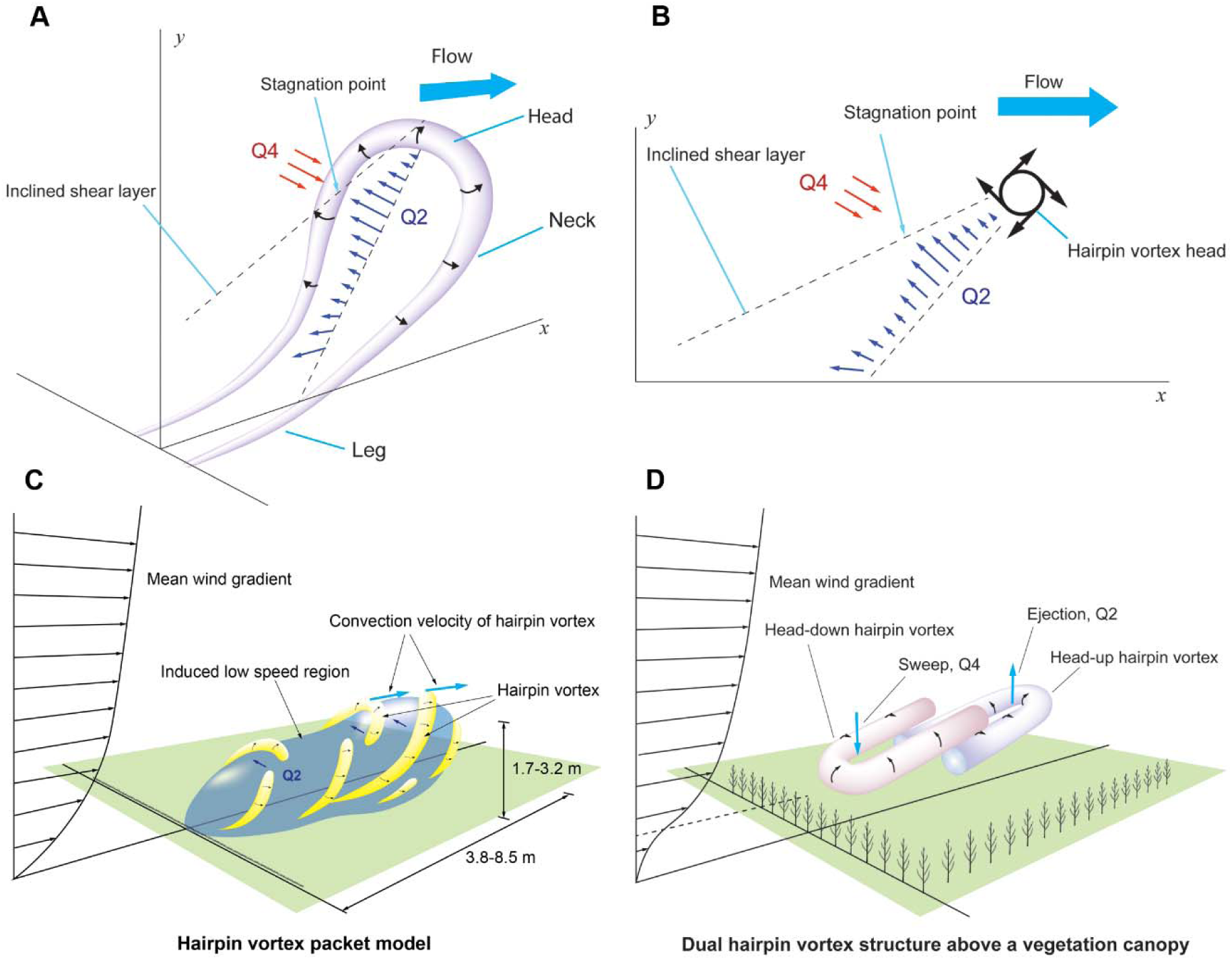
(A) Schematic diagram of a single hairpin vortex in the wall boundary layer. Q2 is an “ejection” region whose velocity vectors are u’ < 0 and v’ > 0. Q4 is a “sweep” region whose velocity vectors are u’ > 0 and v’ < 0. (B) Cross-section of the x-y plane of the hairpin vortex. (C) Schematic diagram of the hairpin vortex packet. Yellow colors mean hairpins or cane-type vortices. Blue region means low momentum region, which contains upward air currents. (D) Coherent structure, “dual hairpin vortex,” on the plant field. Head-down hairpin vortex produces “sweep” event. Head-up hairpin vortex produces “ejection” event. (A, B) Redrawn from [65] (C) Redrawn from [65, 66] (D) Redrawn from [67]

### Shear flow and the spider’s posture

While Reynolds postulated the tangled shape of ballooning fibers in his simulation, the diagonally lying shape of the spider’s ballooning fibers was observed in our field observations in the Teltow canal (see Fig. 7) [16]. The difference between our observation and Reynolds’ simulation is caused by the fact that while Reynolds introduced homogeneous turbulence in his simulation, the real turbulence near the surface boundary layer includes instantaneous horizontal wind shears that are mostly induced by vertical differences in wind speed [68,69]. This diagonally stretched shape of ballooning fibers may be caused by the shear wind in the atmospheric boundary layer. We think that this may be helpful for long-endurance ballooning flight, because horizontally stretched silks produce more drag, up to a factor of 2, than vertically distributed shapes of silk, because of an anisotropic drag of silks in a low Reynolds number flow [70] (see S9).

The observed facts, that spiders outstretch all legs outwards during their flight, is puzzling because of its small drag ratio compared to the drag of a spider’s ballooning silks. Suter concluded that when a spider uses a relatively short length of silk, the influence of posture on its terminal speed is greater than when the silk is very long [19]. However, our observation shows that spiders spin abundant ballooning fibres. From Suter’s empirical formula, spiders’ (weighing 10 to 25 mg) body drags per unit velocity, which are 10.4–14.5 μN · *m*^−1^ · *s*, are just 0.4–0.56% of the whole drag (silks + body) per unit velocity, 2576.5–2580.5 *μ*N *· m*^−1^ · *s*. Even if a spider stretches their legs outwards, the percentage of body drag is still 2.0–2.8% (5-fold change is applied, from Suter’s research) [18,19]. Despite such a small drag effect of spiders’ posture, spiders stretch their legs outwards during flight (see S6C, D). Then the question arises, what role do the stretched legs play in ballooning flight? These should be studied as a future work.

## 5. Conclusion and outlook

Studying ballooning mechanics in spiders can be helpful for understanding not only ecological influence of spiders’ dispersal but also efficient passive transport of particles using air or oceanic currents.

By observing the ballooning behavior in relatively large spiders (10–25 mg *Xysticus* species) in the field and in the laboratory, we have revealed that the large spiders ballooning, even that of 80–150 mg *Stegodyphus* species, is possible with the help of tens of multiple nanoscale fibers. The observed ballooning lines were not identical to the drag line, which has been regarded as a ballooning line. From the observation of behaviors and morphological dimensions of fibers, we concluded that these ballooning silks are two of minor ampullate silks and multiple aciniform silks, which are usually used in other species as wrapping silks. *Xysticus* species, however, used these silks for their aerial dispersal. Spiders showed also interesting behaviors, active sensing motion, such as evaluating the wind conditions before their ballooning behavior. This behavior may save spider’s silk dopes, which can be consumed during their takeoff trials.

Two major features in the physics of ballooning are suggested from the study. First, atmospheric shear flow may be helpful for the high buoyant capability of a ballooning structure, because horizontally/diagonally stretched silks produce more drag than vertically distributed shapes of silks. Second, spiders may use the updrafts that are induced by “coherent structures” in the turbulent atmospheric boundary layer. From the measured wind data, we showed that these updrafts are correlated with lower wind speeds. Therefore, this hypothesis is expected that can explain the fact why spiders usually balloon when the wind speed is lower than 3 m s^-1^. However, these suggestions should be studied further for theoretical firmness.

Whether or not vertical wind speed and fluctuation of wind influence on spiders’ evaluation processes for ballooning, or why spiders stretch their legs outwards during their flight, are questions that still remain. These could be interesting topics for future research.

## 6. Supporting information

**S1 Fig. Ballooning behavior of a male crab spider. The spider shows the tiptoe behavior and spins ballooning silks.**

**S2 Fig. Sequential sketches of ‘tiptoe’ takeoff in the field experiment. Red lines: An anchored line (a drag line). Blue lines: Ballooning lines. (A) A spider, *Xysticus* species, tries to hold the anchored line with one of legs IV. (B) The spider holds the anchored line and then puts it on the substrate. (C) The spider first spins a single or a few number of fibers. (D) And then, the spider first spins many split fibers. (E) If the wind condition is appropriate, the spider releases the substrate. (F) In very short time, the anchored line is cut. The crab spider become airborne.**

**S3 Fig. Sequential sketches of rafting takeoff in the field experiment. Red lines: An anchored line (a drag line). Blue lines: Ballooning lines. (A)(B) A spider, *Xysticus* species, drops down about 0.4 to 1.1 m relying on its anchored line (a drag line) (C) The spider spins a single or a few number of fibers first downstream of the wind. (D) And then, the spider spins many fibers continuously. (E)(F) The spun ballooning lines slowly curved upwards. At some point, the anchored line near the spinnerets is cut and the spider balloons.**

**S4 Fig. The sketch describes the position of the hair dryer. The mixed air zone generates 30°–35°C warm air and fluctuating updraft (horizontal mean wind speed: 0.58 m s**^**-1**^, **vertical mean wind speed: 0.40 m s**^**-1**^).

**S5 Fig. Tidying-up motion of safety line (anchored line) with leg IV. Two independent events; a front view (A), a side view (B). Both spiders hold their safety line and then put it on the substrate before spinning of ballooning lines.**

**S6 Fig. (A, B) An anchored line was found during a tiptoe takeoff. As soon as spiders were airborne, they stretched the legs outwards. (C) To ensure the behavior of outstretched legs during flight, the pose of a spider was observed during its gliding phase. (D) The spider kept its legs outstretched.**

**S7 Table. Numbers of behaviors on the artificial platform.**

**S8 Fig. (A) Spinning motion of ballooning silks in front of the open jet wind tunnel. Ballooning lines were spun from either or both of the median or/and posterior spinnerets. (B) Spinnerets of *Xysticus cristatus* through FESEM. (C) Median spinneret of Xysticus species. Two minor ampullate glands and 10 aciniform glands are distributed on the median spinneret. (D) A number of aciniform glands on the posterior spinneret.**

**S9 Fig. (A) Ballooning structure in a shear flow. (B) Drift of a ballooning structure along with wind. The upper and lower parts of the silk are exposed to the flow fields, which exert to the other directions. (C) The ballooning structure exposed in a shear flow is stretched horizontally. If the drag of a body increases, the structure is stretched more horizontally. (***V*_*wG*_**: Wind speed profile relative to ground**, *V*_*D*_**: Drift speed of a ballooning structure**, *V*_*wR*_**: Wind speed profile relative to a ballooning structure**, *D*_*S*_**: Horizontal component of drag on the spider’s body**, *h* **: height)**

**S10 Movie. A pre-ballooning behavior of a crab spider, *Xysticus* species.**

## 7. Acknowledgements

We would like to thank Jason Dunlop of the Naturkundemuseum Berlin for advice on spider identification. Many thanks to Robert B. Suter for the detailed explanation of his research. We thank Iván Santibáñez Koref and Wladyslaw Chrusciel for the scientific discussions. We also thank Susi Koref, who improved the English in this manuscript. Especially, I would like to acknowledge the state of Berlin (Elsa-Neumann-Scholarship) for the support of PhD scholarship (2015–2018).

## 9. Author contributions

MS.C. designed the study, performed the experiments and wrote the text. I.R. and P.N. supervised the design of the study and contributed to the writing of the text. MS.C and C.F. executed the FESEM observation. All authors gave final approval for publication.

## References

1. Richter CJJ. Aerial dispersal in relation to habitat in eight wolf spider species (Pardosa, Araneae, Lycosidae). Oecologia 1970;5(3):200–14.

2. Salmon JT, Horner NV. Aerial dispersion of spiders in North Central Texas. The J. Arachnol. 1977;5(2):153–7.

3. Tolbert WW. Aerial dispersal behavior of two orb weaving spiders. Psyche: A Journal of Entomology 1977;84(1):13–27.

4. Dean D.A, Sterling WL. Size and phenology of ballooning spiders at two locations in eastern Texas. J. Arachnol. 1985;(13):111–20.

5. Greenstone MH, Morgan CE, Hultsch AL, Farrow RA, Dowse JE. Ballooning spiders in Missouri, USA, and New South Wales, Australia: Family and mass distributions. J. Arachnol. 1987;15(2):163–70.

6. Morse DH. Dispersal of the spiderlings of *Xysticus Emertonu* (Araneae, Thomisidae), a litter-dwelling crab spider. J. Arachnol. 1992;(20):217–221.

7. Cbosby CR, Crosby, C. R. and Bishop, S. C. Aeronautic spiders with a description of a new species. Journal of the New York Entomological Society 1936;1(44):43–9. Available from: http://www.jstor.org/stable/25004641.

8. McCook HC. American spiders and their spinning work: A natural history of the orbweaving spiders of the United States: with special regard to their industry and habits. Landisville, PA: Coachwhip Publications;2006.

9. Blackwall J. Observations and experiments, made with a view to ascertain the means by which the spiders that produce gossamer effect their aerial excursions. Transactions of the Linnean Society of London 1827;(15):449–59.

10. Bell JR, Bohan DA, Shaw EM, Weyman GS. Ballooning dispersal using silk: World fauna, phylogenies, genetics and models. BER 2005;95(02):2.

11. Darwin C. Journal of researches into the natural history and geology of the countries visited during the voyage of H.M.S. Beagle round the world, under the Command of Capt. Fitz Roy, R.N. 2nd ed. New York: John Murray, Albemarle Street;1845.

12. Glick PA. The distribution of insects, spiders, and mites in the air. USDA Tech Bull 1939;(673):1–150.

13. Bilsing SW. Quantitative studies in the food of spiders. The Ohio Journal of Science 1920;20(7):215–60.

14. Bristowe WS. A Preliminary Note on the Spiders of Krakatau. Proceedings of the Zoological Society of London 1931;101(4):1387–400.

15. Hormiga G. Orsonwells, A new genus of giant linyphild spiders (Araneae) from the Hawaiian Islands. Invert. Systematics 2002;16(3):369–448.

16. Reynolds AM, Bohan DA, Bell JR. Ballooning dispersal in arthropod taxa with convergent behaviours: dynamic properties of ballooning silk in turbulent flows. Biol Lett 2006;2(3):371–3.

17. Humphrey JAC. Fluid mechanic constraints on spider ballooning. Oecologia 1987;73(3):469–77.

18. Suter RB. Ballooning in spiders: Results of wind tunnel experiments. Ethology Ecology & Evolution 1991;3(1):13–25.

19. Suter RB. Ballooning: Data from spiders in freefall indicate the importance of posture. J. Arachnol. 1992;2(20):107–13. Available from: http://www.jstor.org/stable/3705774.

20. Suter RB. An aerial lottery: The physics of ballooning in a chaotic atmosphere J. Arachnol. 1999;1(27):281–93. Available from: http://www.jstor.org/stable/3705999.

21. Bell JR, Bohan DA, Le Fevre R, Weyman GS. Can simple experimental electronics simulate the dispersal phase of spider ballooners? J. Arachnol. 2005; 33(2):523–32.

22. Gorham PW. Ballooning Spiders: The Case for Electrostatic Flight 2013. Available from: https://arxiv.org/abs/1309.4731v2.

23. Sheldon KS, Zhao L, Chuang A, Panayotova IN, Miller LA, Bourouiba L. Revisiting the physics of spider ballooning. Women in Mathematical Biology 2017;8:125–39.

24. Weyman GS. A review of the possible causative factors and significance of ballooning in spiders. Ethology Ecology & Evolution 1993; 5(3):279–91.

25. Wickler W, Seibt U. Aerial dispersal by ballooning in adult *Stegodyphus mimosarum*. Naturwissenschaften 1986;73(10):628–9.

26. Schneider JM, Roos J, Lubin Y, Henschel JR. Dispersal of Stegodyphus Dumicola (Araneae, Eresidae): They Do balloon after all! J. Arachnol. 2001;(29):114–6.

27. Simonneau M, Courtial C, Pétillon J. Phenological and meteorological determinants of spider ballooning in an agricultural landscape. C R Biol 2016; 339(9-10):08–16.

28. Coyle FA, Greenstone MH, Hultsch AL, Morgan CE. Ballooning mygalomorphs: estimates of the masses of Sphodros and ummidia ballooners (Araneae: Atypidae, Ctenizidae). J. Arachnol. 1985; 13(3):291–6.

29. Greenstone MH, Morgan CE, Hultsch AL. Spider ballooning: development and evaluation of field trapping methods. J. Arachnol. 1985;13(3):337–45.

30. Greenstone MH. Meteorological determinants of spider ballooning: the roles of thermals vs. the vertical windspeed gradient in becoming airborne. Oecologia 1990;84(2):164–8.

31. Vugts HF, Van Wingerden WKRE. Meteorological aspects of aeronautic behaviour of spiders. Oikos 1976;27(3):433.

32. Bristowe WS. The comity of spiders. Lymington: Pisces Conservation/Ray Society;2004.

33. Duffey E. Aerial dispersal in a known spider population. The Journal of Animal Ecology 1956;1(25):85.

34. Bishop L. Meteorological aspects of spider ballooning. Environmental Entomology 1990;19(5):1381–7.

35. Van Wingerden WKRE, Vugts, HF. Factors influencing aeronautic behaviour of spiders. Bulletin of the British Arachnological Society 1974;3(1):6–10

36. Richter C. Some aspects of aerial dispersal in different populations of wolf spiders, with particular reference to Pardosa Amentata (Araneae Lycosidae). Miscellaneous Papers, Landbouwhogeschool, Wageningen 1971;(8):77–88.

37. Lee VMJ, Kuntner M, Li D. Ballooning behavior in the golden orb web spider Nephila pilipes (Araneae: Nephilidae). Front. Ecol. Evol. 2015;3:e86780.

38. Zhao L, Panayotova IN, Chuang A, Sheldon KS, Bourouiba L, Miller LA. Flying spiders: Simulating and modeling the dynamics of ballooning. Women in Mathematical Biology 2017;8:179–210.

39. Henschel JR. Dispersal mechanisms of Stegodyphus (Eresidae): Do they balloon? J. Arachnol. 1995;(23):202–4.

40. Willmarth WW, Lu SS. Structure of the Reynolds stress near the wall. J. Fluid Mech. 1972;55(01):65.

41. Lu SS, Willmarth WW. Measurements of the structure of the Reynolds stress in a turbulent boundary layer. J. Fluid Mech. 1973;60(03):481.

42. Adrian RJ. Hairpin vortex organization in wall turbulence. Physics of Fluids 2007;19(4):041301.

43. Zhu W, van Hout R, Katz J. PIV Measurements in the atmospheric boundary layer within and above a mature corn canopy. Part II: Quadrant-hole analysis. J. Atmos. Sci. 2007;64(8):2825–38.

44. Steiner AL, Pressley SN, Botros A, Jones E, Chung SH, Edburg SL. Analysis of coherent structures and atmosphere-canopy coupling strength during the CABINEX field campaign. Atmos. Chem. Phys. 2011;11(23):11921–36.

45. Palmgren P. Experimentelle Untersuchungen Fiber die Funktion der Trichobothrien bei Tegenaria Derhami. Soc. Acta Zool. Fenn. 1936;(19):3–28.

46. Washburn JO, Washburn L. Active aerial dispersal of minute wingless arthropods: exploitation of boundary-layer velocity gradients. Science 1984;223(4640):1088–9.

47. Hunt JC, Morrison JF. Eddy structure in turbulent boundary layers. European Journal of Mechanics - B/Fluids 2000;19(5):673–94.

48. McNaughton KG. Turbulence structure of the unstable atmospheric surface layer and transition to the outer layer. Boundary-Layer Meteorology 2004;112(2):199–221.

49. Barth F, Komarek S, Humphrey J, Treidler B. Drop and swing dispersal behavior of a tropical wandering spider: Experiments and numerical model. J Comp Physiol A 1991;169(3).

50. Coyle FA. Aerial dispersal by mygalomorph spiderlings (Araneae, Mygalomorphae). J. Arachnol. 1983;(11):283–6.

51. Vollrath F. Strength and structure of spiders’ silks. Journal of Biotechnology 2000; 74(2):67–83.

52. Moon MJ, An JS. Spinneret Microstructure of the Silk Spinning Apparatus in the Crab Spider, Misumenops tricuspidatus (Araneae: Thomisidae). Entomol Research 2005;35(1):67–74.

53. Osaki S. Is the mechanical strength of spider’s drag-lines reasonable as lifeline? International Journal of Biological Macromolecules 1999;24(2–3):283–7.

54. Eberhard WG. How spiders initiate airborne lines. J. Arachnol. 1987;1(15):1–9.

55. Ko FK, Kawabata S, Inoue M, Niwa M, Fossey S, Song JW. Engineering properties of spider silk. MRS Proc. 2001;702:91.

56. Bonino MJ. Material Properties of Spider Silk. dissertation. Rochester, NY: The College School of Engineering and Applied Sciences;2003.

57. Griswold CE. Atlas of phylogenetic data for entelegyne spiders (Araneae: Araneomorphae: Entelegynae): With comments on their phylogeny. In: Proceedings of the California Academy of Sciences, 4th series (Vol. 56, Suppl. 2) San Francisco: California Academy of Sciences;2005.

58. Purcell EM. Life at low Reynolds number. Am. J. Phys. 1977;45:3–11.

59. Roos FW, Willmarth WW. Some experimental results on sphere and disk drag. AIAA Journal 1971;9(2):285–91.

60. Pelegrini MF, Vieira EDR. Flow past a sphere moderate Reynolds numbers. 17th International Congress of Mechanical Engineering, November 10–14, 2003, São Paulo, SP.

61. Dudley R. The biomechanics of insect flight: Form, function, evolution. Princeton, NJ: Princeton Univ. Press;2002.

62. Sane SP. The aerodynamics of insect flight. Journal of Experimental Biology 2003;206(23):4191–208.

63. Bomphrey RJ, Nakata T, Phillips N, Walker SM. Smart wing rotation and trailing-edge vortices enable high frequency mosquito flight. Nature 2017;544(7648):92–5.

64. Theodorsen T. Mechanism of turbulence. In Proc. Midwestern Conf. Fluid Dyn., Ohio State University, Columbus, Ohio. 1952.

65. Adrian RJ, Meinhart CD, Tomkins CD. Vortex organization in the outer region of the turbulent boundary layer. J. Fluid Mech. 2000;422:1–54.

66. Hommema SE, Adrian RJ. Packet structure of surface eddies in the atmospheric boundary layer. Boundary-Layer Meteorology 2003;106(1):147–70.

67. Finnigan JJ, Shaw RH, Patton EG. Turbulence structure above a vegetation canopy. J. Fluid Mech. 2009;637:387.

68. Dennis DJC. Coherent structures in wall-bounded turbulence. An Acad Bras Cienc 2015;87(2):1161–93.

69. Grass AJ. Structural features of turbulent flow over smooth and rough boundaries. J. Fluid Mech. 1971;50:233.

70. Childress S. Mechanics of swimming and flying. In: Cambridge studies in mathematical biology, vol. 2.: Cambridge University Press;1981. Available from: https://doi.org/10.1017/CBO9780511569593.

