## Supplementary figures and images for "An Observational Study of Ballooning in Large Spiders: Nanoscale Multi-Fibers Enable Large Spiders’ Soaring Flight"

### Supplementary Materials

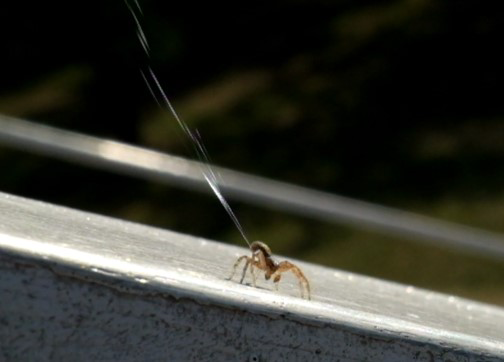

### Supplementary Materials

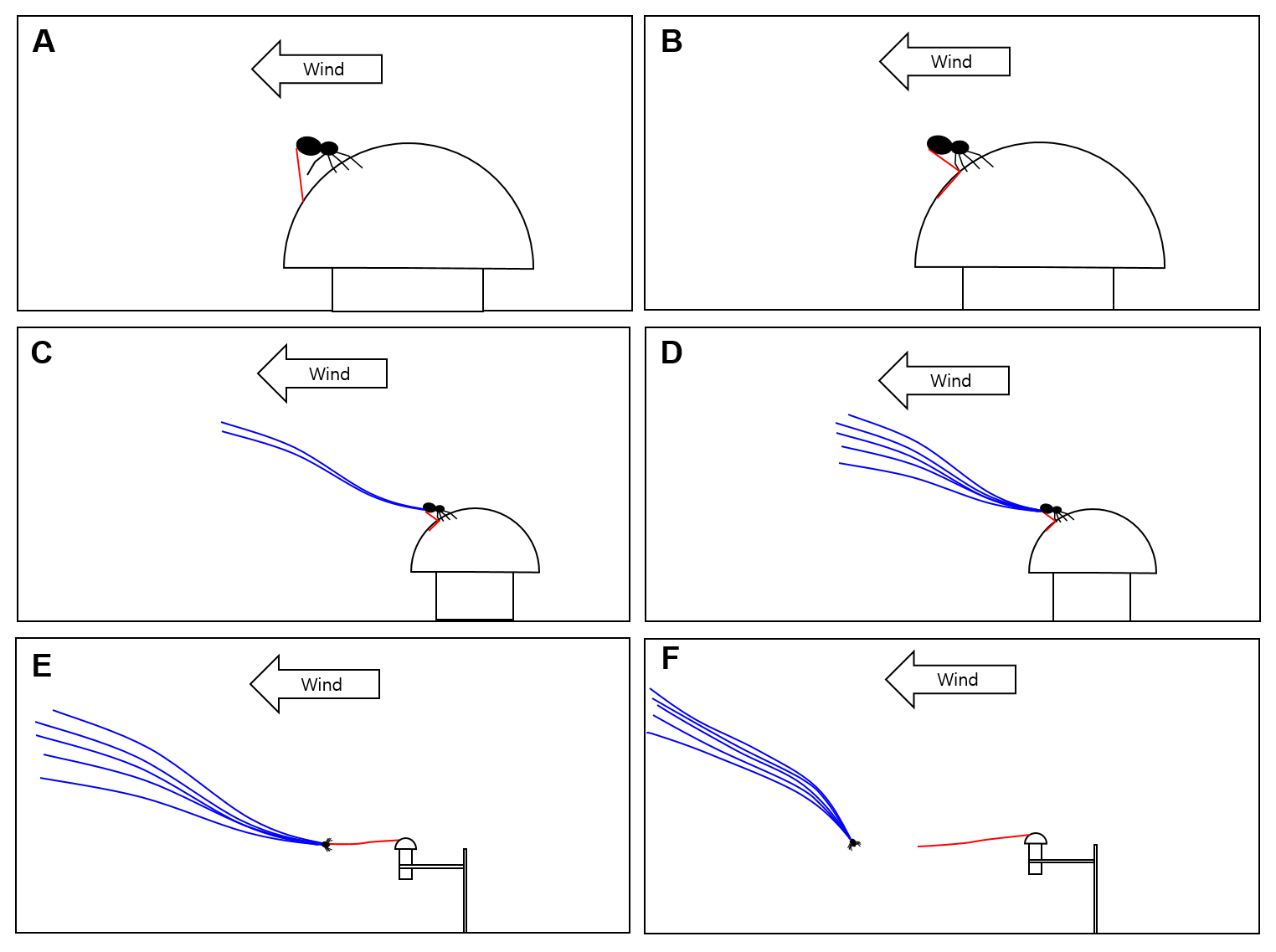

### Supplementary Materials

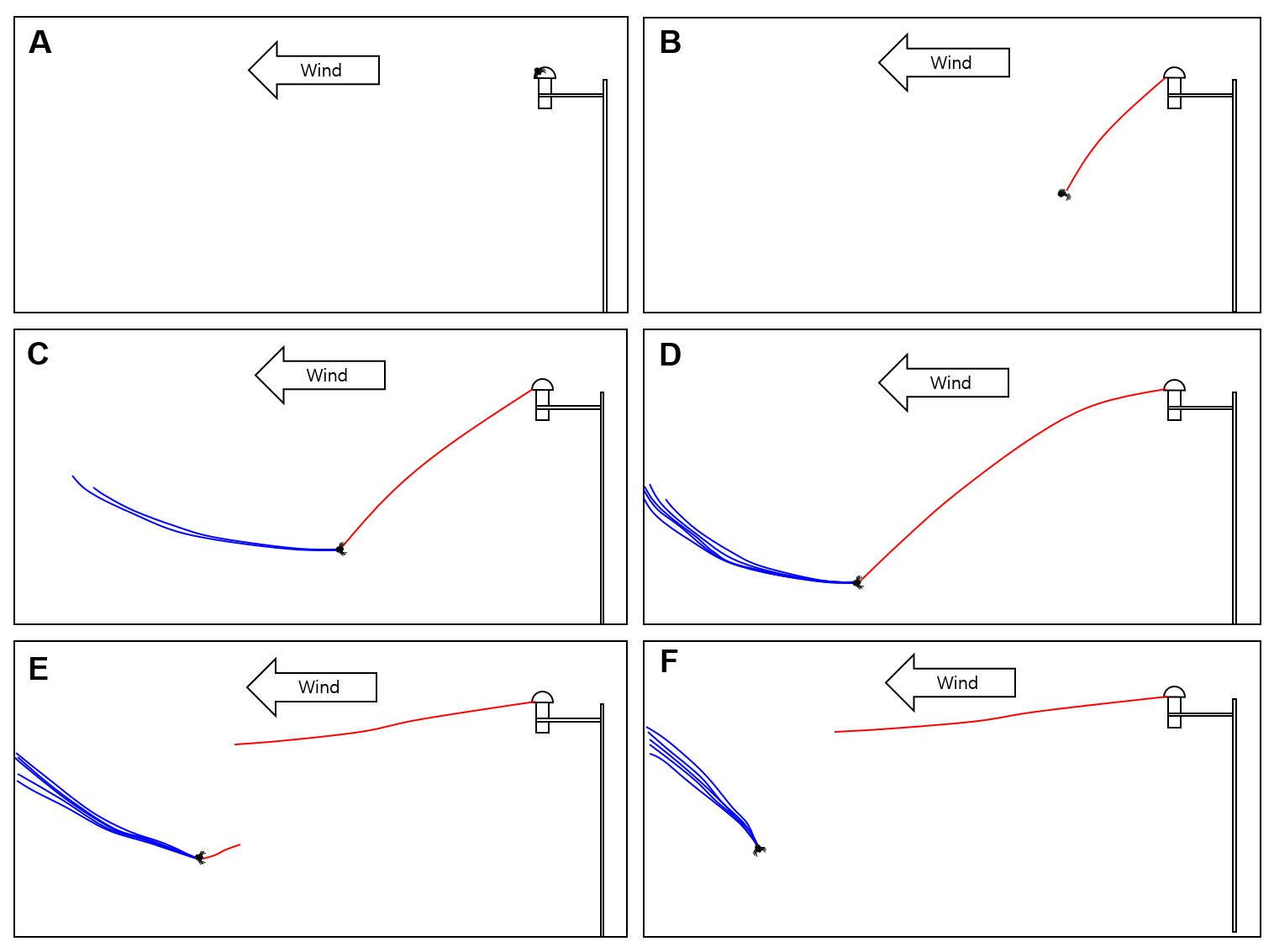

### Supplementary Materials

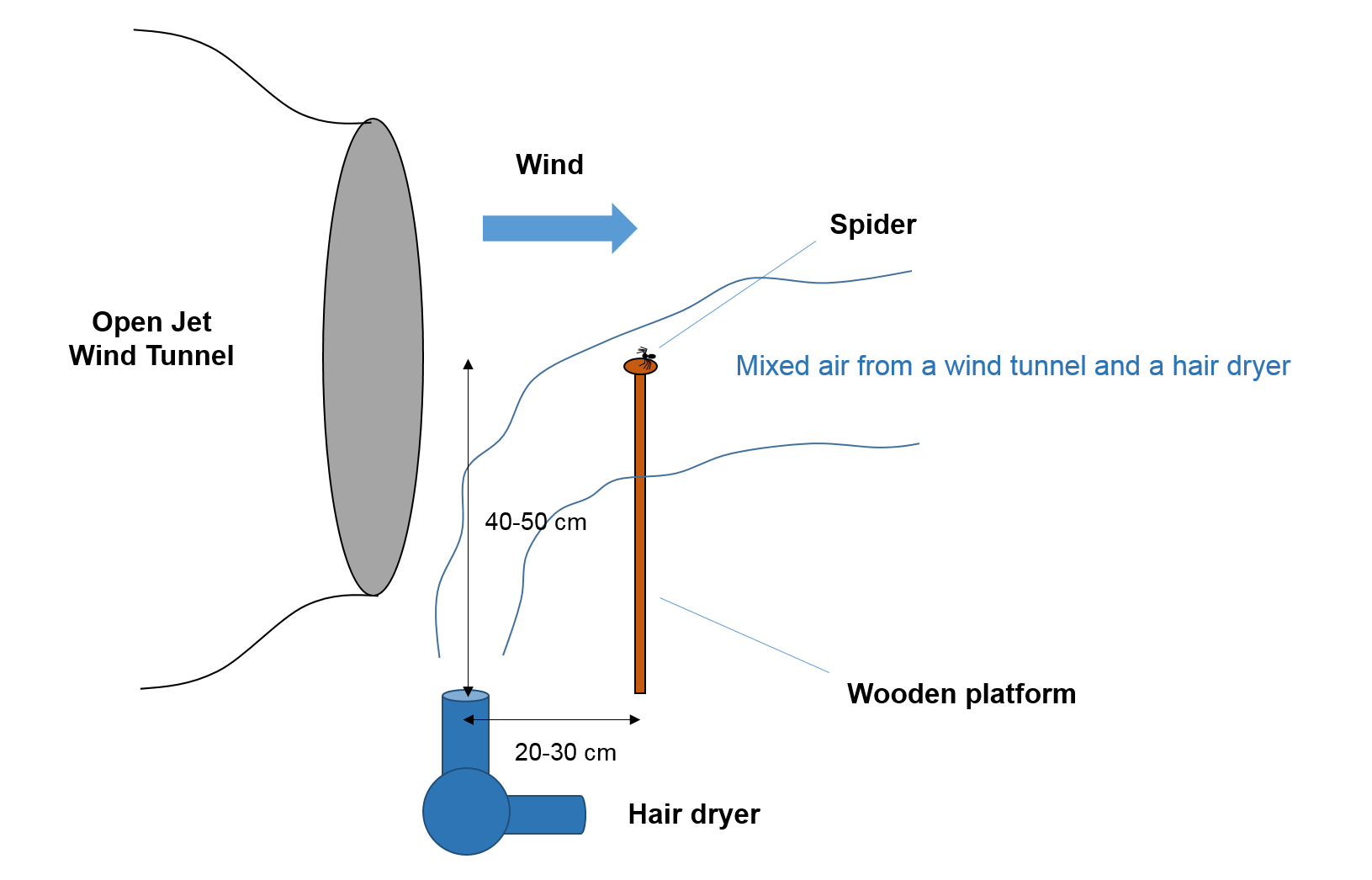

### Supplementary Materials

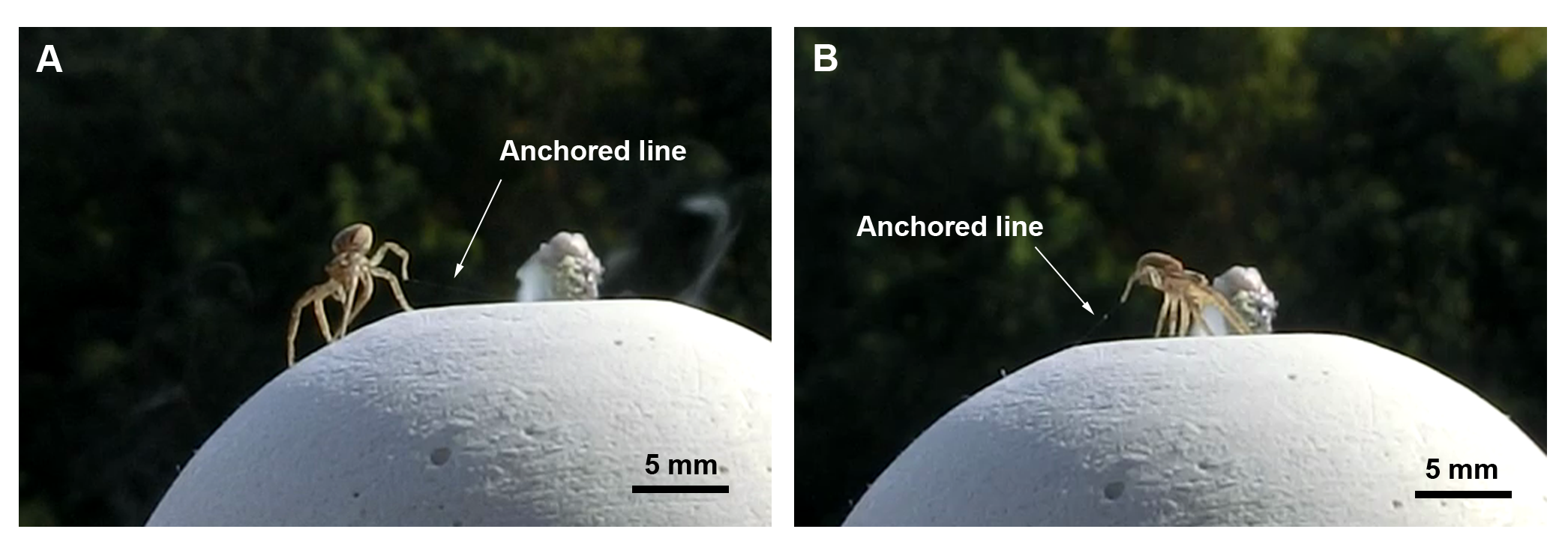

### Supplementary Materials

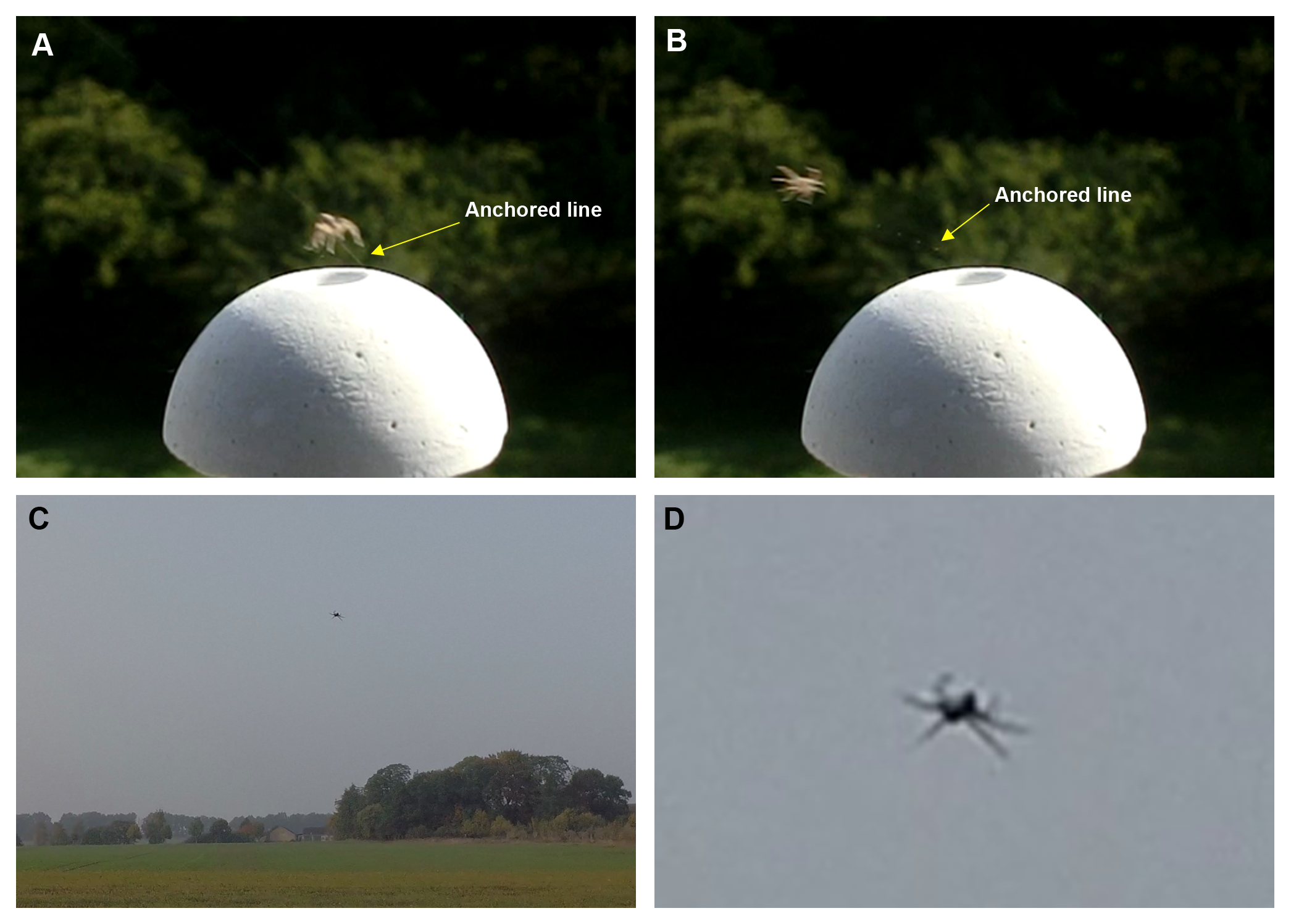

### Supplementary Materials

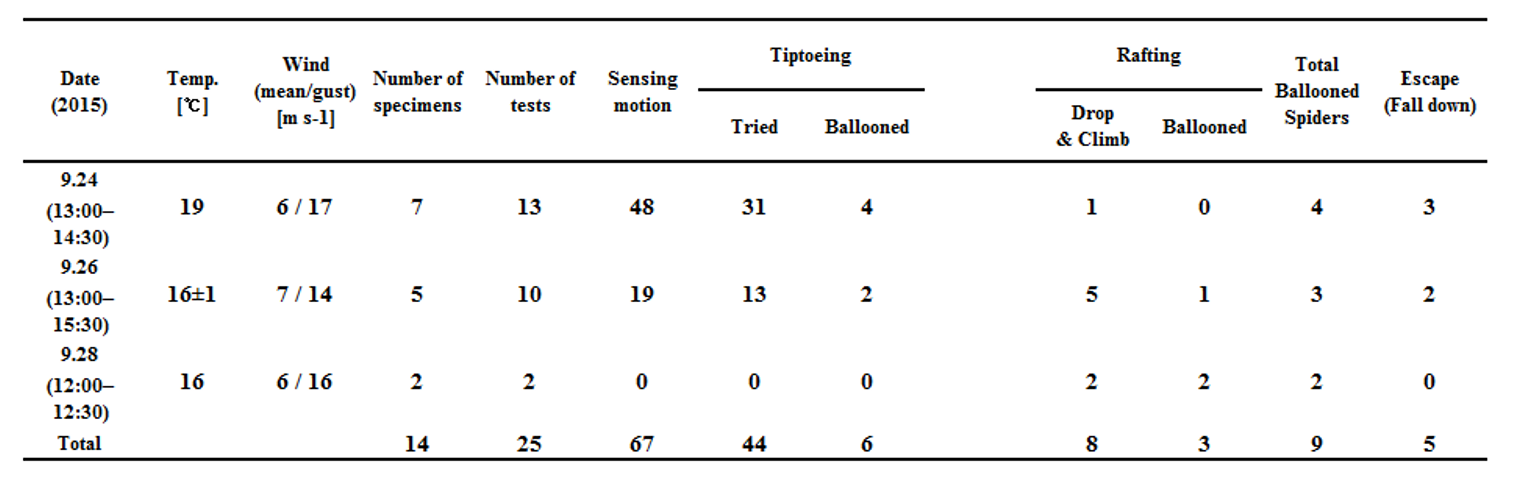

### Supplementary Materials

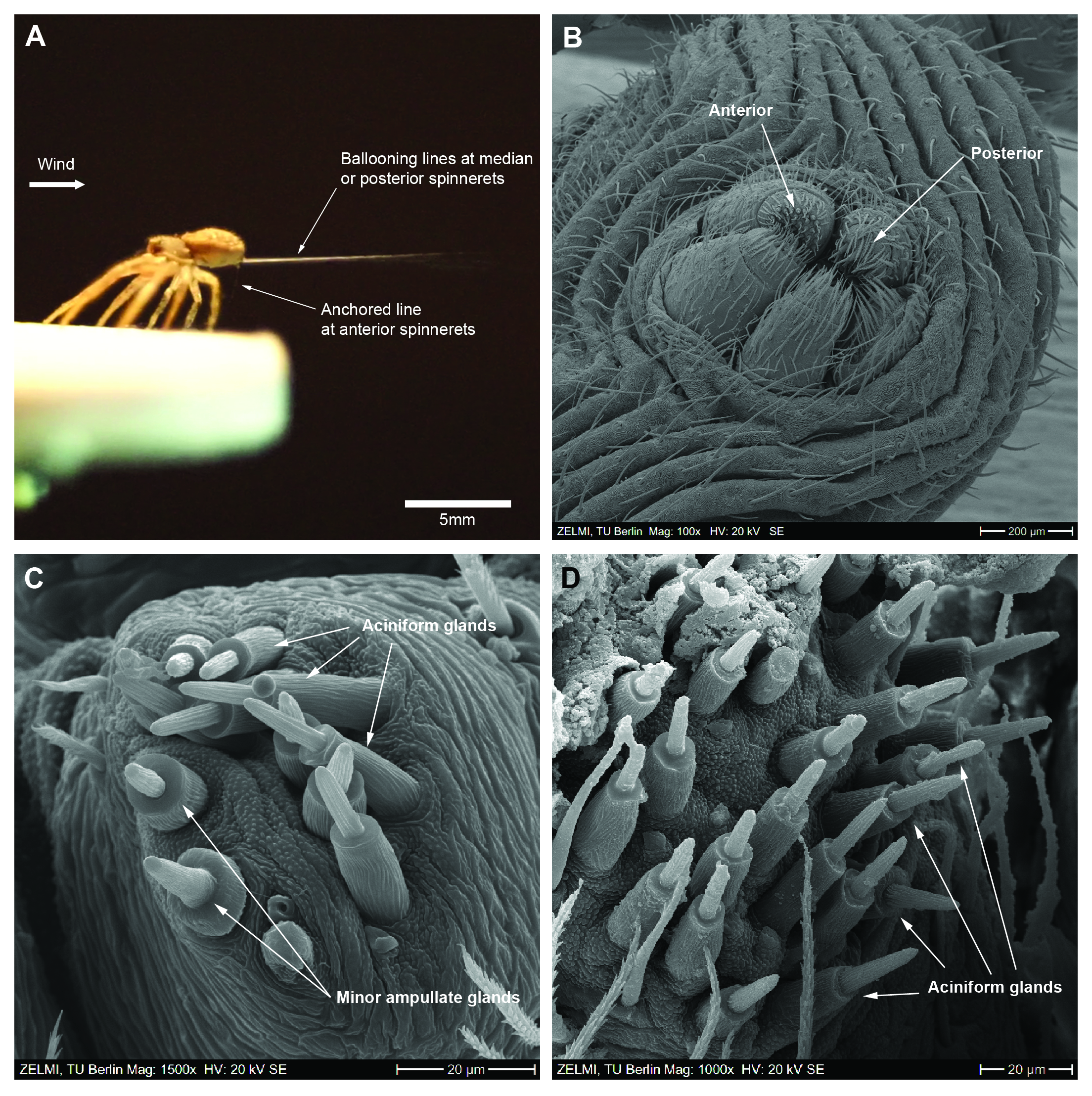

### Supplementary Materials

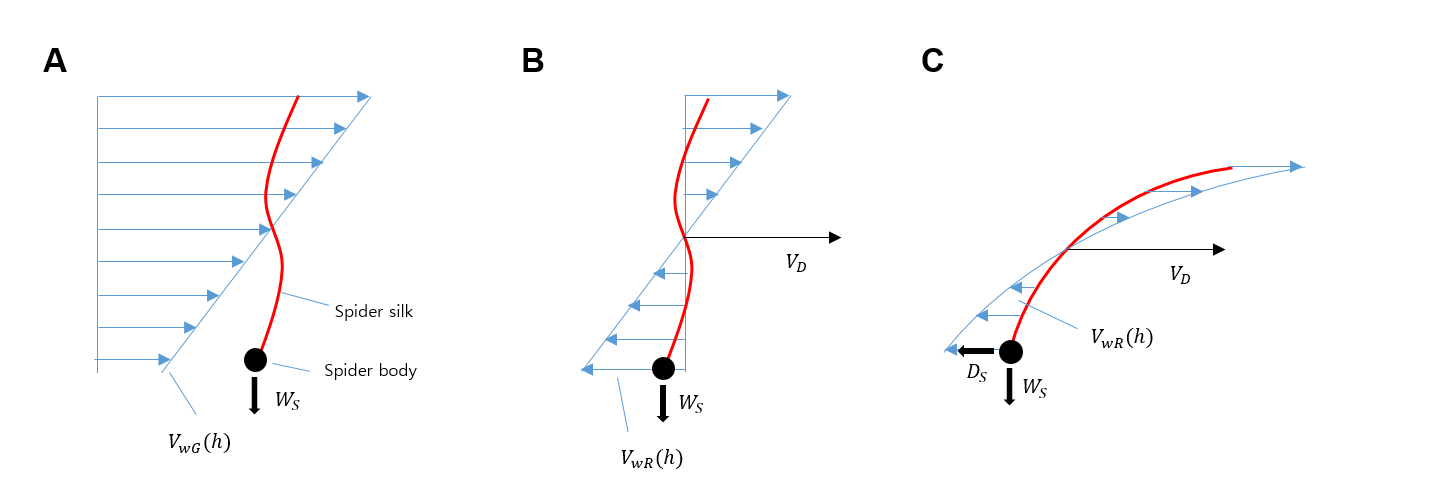
